# TCR-pMHC bond length controls TCR ligand discrimination

**DOI:** 10.1101/433938

**Authors:** Dibyendu K. Sasmal, Wei Feng, Sobhan Roy, Peter Leung, Yanran He, Chufan Cai, Guoshuai Cao, Huada Lian, Jian Qin, Enfu Hui, Hans Schreiber, Erin Adams, Jun Huang

## Abstract

T-cell receptors (TCRs) detect specifically and sensitively a small number of agonist peptide-major histocompatibility complexes (pMHCs) from an ocean of structurally similar self-pMHCs to trigger antigen-specific adaptive immune responses^1–4^. Despite intense efforts, the mechanism underlying TCR ligand discrimination remains a major unanswered question in immunology. Here we show that a TCR discriminates between closely related peptides by forming TCR-pMHC bonds with different lengths, which precisely control the accessibility of CD3ζ immunoreceptor tyrosine-based activation motifs (ITAMs) for phosphorylation. Using *in situ* fluorescence resonance energy transfer (FRET)^3,5^, we measured the intermolecular length of single TCR-pMHC bonds and the intramolecular distance of individual TCR-CD3ζ complexes at the membrane of live primary T cells. We found that an agonist forms a short TCR-pMHC bond to pull the otherwise sequestered CD3ζ off the inner leaflet of the plasma membrane, leading to full exposure of its ITAMs for strong phosphorylation. By contrast, a structurally similar weaker peptide forms a longer bond with the TCR, resulting in partial dissociation of CD3ζ from the membrane and weak phosphorylation. Furthermore, we found that TCR-pMHC bond length determines 2D TCR binding kinetics and affinity, T-cell calcium signaling and T-cell proliferation, governing the entire process of signal reception, transduction and regulation. Thus, our data reveal the fundamental mechanism by which a TCR deciphers the structural differences between foreign antigens and self-peptides via TCR-pMHC bond length to initiate different TCR signaling for ligand discrimination.

---

It remains elusive how a TCR deciphers the structural differences among peptides and properly propagates surface recognition signals across the cell plasma membrane to CD3 intracellular ITAM domains to induce distinct T-cell signaling. The TCR conformational change model postulates a conformational change in a TCR upon pMHC binding, but no conformational changes at the binding interface have been identified that are conserved in TCR-pMHC crystal structures^6,7^. However, crystal structures only provide a “snapshot” of the thermodynamically stable conformation of purified TCR and pMHC proteins. To carefully examine possible TCR conformational changes and critically probe the mechanisms of TCR ligand discrimination *in situ*, we used FRET^5,8^, a spectroscopic ruler with subnanometer precision, to measure the intermolecular length of a TCR-pMHC bond and the intramolecular distance of a TCR-CD3ζ complex at the immunological synapse of a live primary 5C.C7 transgenic CD4^+^ T cell in real time with a temporal resolution of ~10 ms.

To measure the intermolecular length of a single TCR-pMHC bond, a TCR and a pMHC were site-specifically labeled with a FRET acceptor Cy5 and a FRET donor Cy3, respectively^3^. Because the TCR was labeled via an anti-TCR single-chain variable fragment (scFv) J1 (Fig. 1a and Extended Data Fig. 1), the distance between the Cy3 and the Cy5 is a pseudo TCR-pMHC bond length. Yet, it nevertheless provides a reasonable approximation of the length of a single TCR-pMHC bond. The intermolecular length of a single TCR-pMHC bond on the cell surface was measured by Cy3/Cy5 FRET (FRET1) in real-time (Fig. 1a). To measure the intramolecular distance between the TCR and the CD3ζ in a transmembrane TCR-CD3ζ complex, we inserted a green fluorescent protein (GFP) to the C-terminus of the CD3ζ chain, and the TCR was labeled with an Alexa568 fluorophore using a different anti-TCR scFv J3 with a unique labeling site close to the cell membrane^3^. The real-time intramolecular distances of TCR-CD3ζ complexes were measured by GFP/Alexa568 FRET (FRET2) using time-lapse microscopy (Fig. 1a and Extended Data Fig. 2a). We first performed experiments to assess the feasibility and specificity of the cell surface FRET1 and transmembrane FRET2 on lipid bilayer and glass surface containing pMHCs and accessary molecules ICAM-1 and B7-1, respectively (Fig. 1b-e, Extended Data Fig. 2a and 3-4, Supplimentary Movie 1-3). For cell surface FRET1, we readily detected FRET signals for three agonist pMHCs but not for a null pMHC on lipid bilayer and glass surface, and the FRET efficiencies (E_FRET_) were positively correlated with the pMHC potencies in activating T cells^9^. The average synaptic E_FRET1_ was 0.79, 0.54 and 0.29 for the super agonist K5, agonist MCC and weak agonist 102S, respectively (Fig. 1d). However, no synaptic FRET was found for the null pMHC (Fig. 1b and d). These data validated the specificity of our cell surface TCR-pMHC FRET and were consistent with a previous report^3^. In contrast, the transmembrane TCR-CD3ζ FRET efficiencies were inversely correlated with the pMHC potencies with the highest FRET observed for the null ligand (Fig. 1c and e). In the presence of agonist pMHCs K5, MCC and 102S, the transmembrane FRET was only detected at the TCR-CD3ζ co-localized microclusters but not outside of the co-localized microclusters (Fig. 1c and Extended Data Fig. 5a). Replacing the FRET acceptor Alexa568 with a Cy5 dye abolished the transmembrane FRET (Extended Data Fig. 5b). These experiments confirmed the specificity of the transmembrane TCR-CD3ζ FRET as well.

**Fig. 1.**
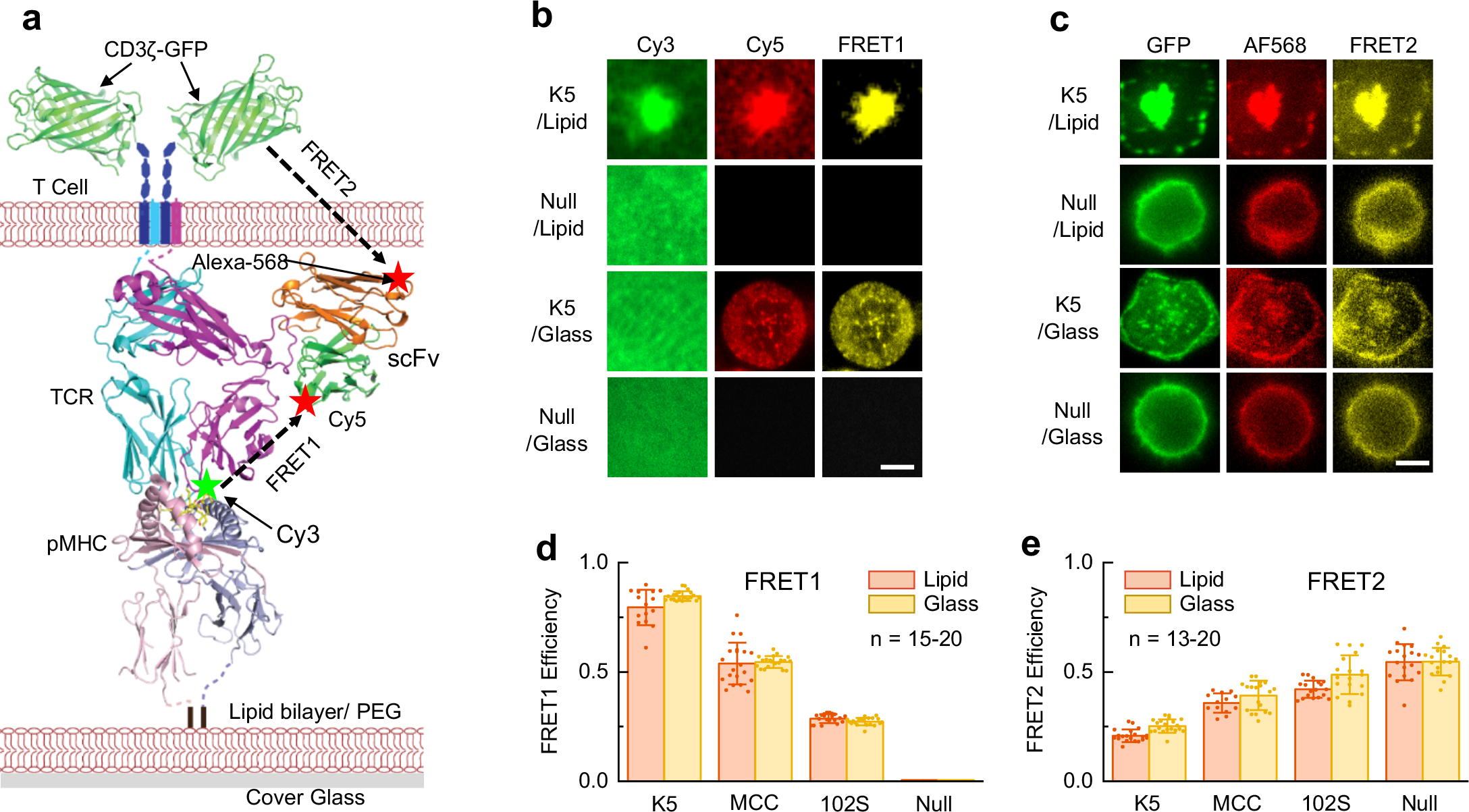
Measurement of TCR conformational dynamics by FRET. **a**, A composite structural model of TCR, pMHC, scFv and CD3ζ-GFP. For measuring TCR-pMHC bond conformational dynamics, the TCR was labeled by a Cy5 via a scFv J1 and the peptide was labeled by a Cy3. For determining TCR-CD3ζ conformational changes, the TCR was labeled by an Alexa568 via a scFv J3 and the CD3ζ was tagged by a GFP. Extracellular Cy3/Cy5 FRET1 and transmembrane GFP/Alexa568 FRET2 were indicated by dashed lines. All pMHCs were anchored on either lipid bilayer or PEG-Ni^2+^ glass surface (Extended Data Fig. 1). Note: J1 and J3 are different scFvs and each only has a unique labeling site. **b-c**, Donor, acceptor and FRET signals of TCRs interacting with K5 or Null pMHC on lipid bilayer or glass surface. Shown were representative data of 3-5 independent experiments for each peptide at 37 °C. Scale bar is 5 μm. **d-e**, FRET efficiencies measured for K5, MCC, 102S and null pMHCs on lipid bilayer (red) and glass surface (yellow). At least 13 cells were used for determining the FRET efficiency for each pMHC. Also see Extended Data Fig. 1–4 and Supplementary Movie 1-3.

**Fig. 2.**
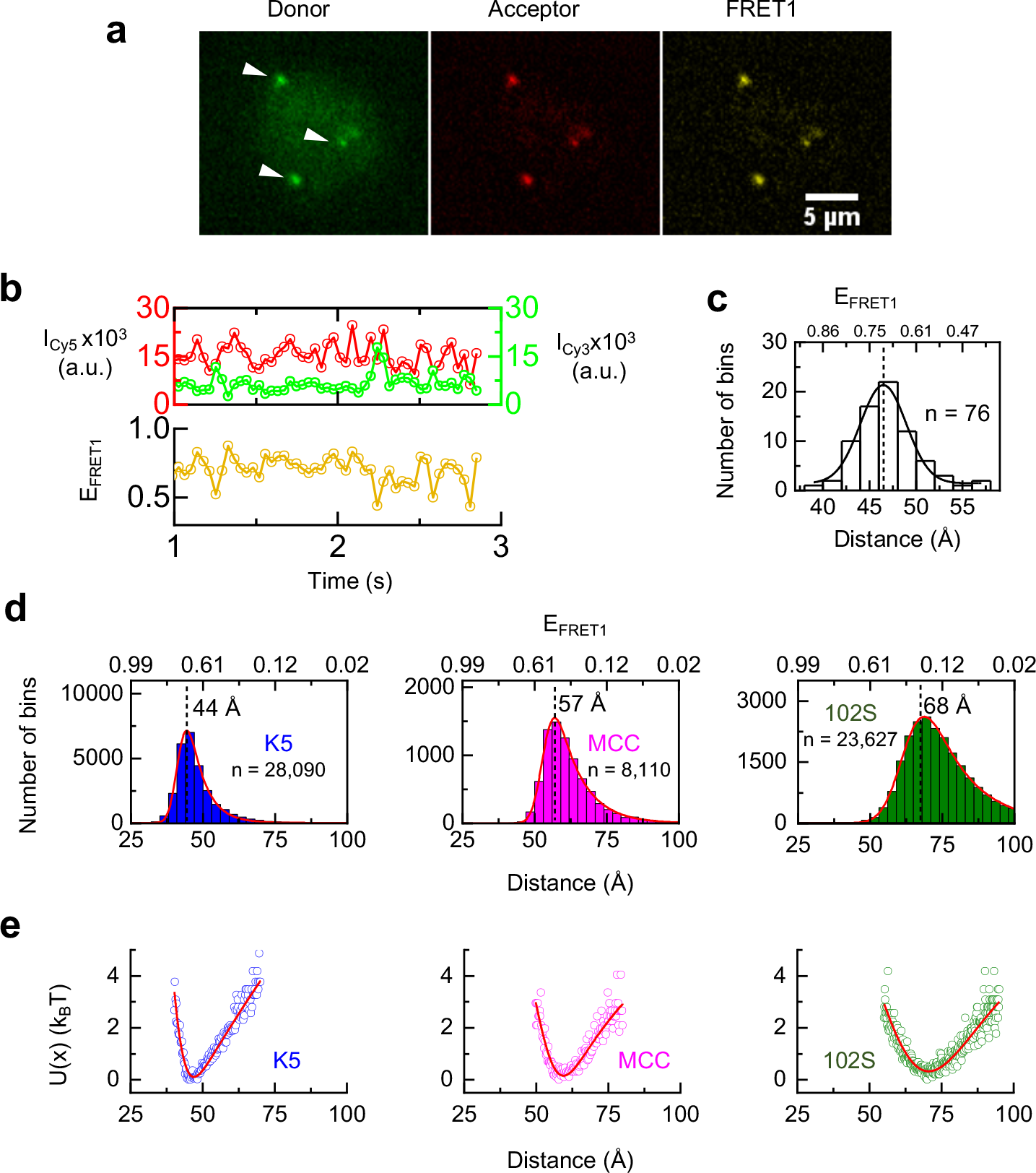
Single TCR-pMHC bond length and conformational dynamics. **a**, A representative smFRET event mediated by the interaction between a TCR and a K5 pMHC. Shown were the donor, acceptor, and FRET signals and white arrows indicated single molecules. **b**, Single-molecule time trajectories of the donor (green, Cy3-pMHC) and the acceptor (red, Cy5-TCR) intensities (upper panel) and the corresponding time trajectory of smFRET efficiency (yellow, lower panel). Also see Extended Data Fig. 7. **c**, Histogram of the Cy3-Cy5 distances (pseudo bond length) calculated from smFRET efficiencies from Fig. 2b (lower panel) and fitted with a Gaussian distribution (black curve). Also see Extended Data Fig. 6a. **d**, Histograms of pseudo TCR-pMHC bond lengths for K5, MCC and 102S pMHCs. Each histogram used 8,000-28,000 bond lengths to identify the most probable bond length for each peptide. Also see Extended Data Fig. 6–8 and Supplementary Movie 4. **e**, Potential-of-mean-force (PMF) of a TCR interacting with K5, MCC and 102S pMHCs, respectively.

We next performed Cy3/Cy5 single-molecule FRET (smFRET) on the lipid bilayer to measure the lengths of single TCR-pMHC bonds using total internal reflection fluorescence (TIRF) microscopy (Fig. 2, Extended Data Fig. 2a and Supplimentary Movie 4). TCR-pMHC bond formation at the live T-cell membrane brought the donor Cy3 and the acceptor Cy5 into close proximity to enable smFRET (Fig. 1a and 2a). The fluorescent intensities of Cy3 and Cy5 were real-time recorded with a rate of ~100-200 frames/s (Fig. 2b, top panel and Extended Data Fig. 6a), and the E_FRET_ values were calculated based on the intensities of the donor Cy3 and the acceptor Cy5 (Methods and Fig. 2b, bottom panel). As E_FRET_ is inversely proportional to the sixth power of the distance between the donor and the acceptor, the Cy3/Cy5 smFRET serves as a sensitive microscopic ruler to precisely measure the TCR-pMHC bond length in real-time. The time trajectory of E_FRET_ showed that a TCR-pMHC bond is dynamic and displays a range of continuous conformations (Fig. 2b), aligned with a recent report that a TCR undergoes conformational changes upon ligation^10^. Recording the conformational fluctuation trajectories provided a real-time observation of single TCR-pMHC bond dynamics. By plotting the E_FRET_ histogram and fitting it with a Gaussian function, we identified the most probable E_FRET_ value of 0.7, corresponding to a 47 Å distance between the 5C.C7 TCR and the super agonist K5 pMHC in a representative smFRET trajectory (Fig. 2c). We repeated our single-bond measurements for K5 pMHC and also performed similar smFRET experiments for agonist MCC and weak agonist 102S pMHCs (Extended Data Fig. 7). After collecting many individual smFRET trajectories for each pMHC, we pooled all E_FRET_ data together and plotted the histograms for each pMHC (≥8,000-28,000 events from 1,300-1,800 trajectories per histogram, Fig. 2d). Remarkably, the distributions of E_FRET_ showed that the pseudo TCR-pMHC bond lengths were peptide dependent, and single TCR-pMHC bonds were highly dynamic with a range of continuous conformational states. Fitting each histogram with a Gaussian function (curves, Fig. 2d) yielded the most probable bond length for each pMHC: 44 Å for K5 (super agonist), 57 Å for MCC (agonist), and 68 Å for 102S (weak agonist) (Fig. 2d), directly revealing angstrom-level, ligation-induced, peptide-dependent TCR-pMHC bond lengths and conformational dynamics *in situ*. This key information has been missing from previously reported TCR-pMHC crystal structures^7,11^. We further measured the average pseudo TCR-pMHC bond lengths using ensemble FRET (Extended Data Fig. 8) on both a lipid bilayer (Fig. 1a) and a glass surface (Extended Data Fig. 1), confirming our single bond measurements obtained using smFRET (Fig. 2d). We then quantified the binding strength of each single TCR-pMHC bond by analyzing the potential-of-mean-force (PMF)^12^, which measures the free energy cost of variation in bond length. PMF minimizes at equilibrium, and its curvature governs the size of fluctuation (Fig. 2e). Clearly, super agonist K5 and agonist MCC have deep and narrow energy wells, indicating strong and stable bonds. In contrast, the weak agonist 102S has a shallow and wide energy well, suggesting weak bond strength and unstable binding state. Overall, the depth and width of the PMF revealed that K5 and MCC form more stable (shorter) bonds with TCRs compared to that of 102S (Fig. 2e), consistent with previous reports that K5 and MCC have higher 3D *in vitro* binding affinities with TCRs than that of 102S^3,9^ as well as that TCR triggering is dependent on receptor-ligand complex dimensions^13,14^. Our measurements not only discovered that TCR triggering is critically dependent on the length of a TCR-pMHC bond but also linked bond length to bond energy, thus providing fundamental knowledge for understanding TCR discrimination and signaling.

It is not clear how CD3ζ dissociates from the membrane to initiate T-cell intracellular signaling. To further understand how surface TCR-pMHC bonds propagate extracellular recognition signals to intracellular CD3ζ ITAMs across the cell membrane, we developed a transmembrane FRET assay to measure the conformational change between the extracellular TCR domain and intracellular CD3ζ chain of the same molecular complex (Fig. 1a and Extended Data Fig. 1). Upon the addition of 5C.C7 transgenic T cells with Alexa568-labeled TCRs and GFP-tagged CD3ζ to the lipid bilayer or glass surface containing pMHC ligands, we observed rapid microcluster formation and instant onset of transmembrane FRET between TCRs and CD3ζ. TCRs were found colocalized with CD3ζ in plasma membrane microclusters [Pearson correlation coefficient (PCC)^15^, 0.93 ± 0.07 for lipid bilayer and 0.59 ± 0.19 for glass surface] (Fig. 3a-b, Extended Data Fig. 6b and 9–10, Supplymentary Movie 5-7), highlighting the initiation of T-cell signaling at the segregated TCR-pMHC bond-mediated close contacts^16^. The high degree of TCR-CD3ζ colocalization suggested obligate assembly of TCR-CD3ζ for effective T-cell signaling and verified the specificity of our transmembrane TCR-CD3ζ FRET. On the lipid bilayer, TCRs and CD3ζ quickly formed relatively large microclusters, which continuously moved from the periphery to the center of the cell and formed the immunological synapse (Fig. 3a). On the glass surface, TCRs and CD3ζ formed smaller microclusters, the size of which increased slightly over time. These microclusters remained at the original sites without forming a synapse (Fig. 3b), consistent with a previous observation in the patterned grids^17^. We then tracked single microclusters and measured their FRET efficiencies individually on both the lipid bilayer and glass surface in real-time. After converting the FRET efficiencies to distances between a TCR and a CD3ζ, we plotted three-dimensional (Fig. 3c-d) figures to simultaneously illustrate the lateral movement of TCR-CD3ζ complexes on lipid bilayer or glass surface (x-y axis) (Extended Data Fig. 11a-b) and the intramolecular TCR-CD3ζ distance changes across the cell membrane (z axis) upon K5 pMHC engagement (Fig. 3c-d). Although the lateral trajectories of TCR-CD3ζ complexes are quite different between lipid bilayer and glass surface due to different anchorages of pMHCs (comparing Fig. 3a-d), their intramolecular TCR-CD3ζ distances within a microcluster at equilibrium phase consistently showed a ~15 Å difference before and after K5 pMHC ligation (Fig. 3c-d). Similarly, we found TCR-CD3ζ conformational changes after TCRs binding to agonist MCC and weak agonist 102S. The extent of conformational change was dependent on the pMHC potency, as MCC and 102S pMHCs triggered ~10 Å and ~5 Å separation between TCR and CD3ζ after TCR-pMHC ligation, respectively. TCR-CD3ζ conformational changes were restricted in TCR/CD3ζ microclusters and no changes were observed outside of the microclusters (Extended Data Fig. 5a and 9-11). These data highlighted that the TCR-CD3ζ conformational changes occurring at the TCR microclusters were driven by TCR-pMHC engagement, consistent with previous reports that TCR microclusters are hotspots and essential structures for TCR signaling^2,18–20^. To better reveal the *in situ* TCR-CD3ζ conformational dynamics, we plotted the time trajectories of intramolecular TCR-CD3ζ distance changes on both the lipid bilayer (Fig. 3e) and glass surface (Fig. 3f) against the stimulation time of each pMHC. We found that the TCR extracellular domain started to separate from its intracellular CD3ζ upon TCR-pMHC ligation, and this process took several minutes to reach a stable and extended TCR-CD3ζ conformational state depending on the pMHC potency, molecular mobility and TCR microcluster size. Interestingly, we found that the TCR-CD3ζ conformational changes have a 5-minute delay on the glass surface compared to those on the lipid bilayer (compare Fig. 3e to 3f), highlighting the importance of molecular mobility and microcluster size in TCR signaling. The delay of TCR-CD3ζ conformational changes was due to the permanent anchorage of pMHCs on glass surface, which decelerated the formation as well as reduced the size of TCR microclusters, the hotspots for signaling. The separation of CD3ζ from the TCR is dependent on the potency of a pMHC, as demonstrated by the amplitude (Fig. 3g) and speed (Fig. 3h) of TCR-CD3ζ conformational changes upon TCR ligation with different pMHCs. Taken with the aforementioned TCR-pMHC bond measurements on the cell surface, our data together showed that the TCR-pMHC bond length directly determines the conformational change of TCR-CD3ζ in an inversely proportional manner. Importantly, our data further showed that TCR-bond length precisely controls the level of CD3ζ phosphorylation through TCR-CD3ζ conformational change (Fig. 3 i-k and Extended Data Fig. 11c). Among three ligands tested, we found that the most potent K5 formed the shortest TCR-pMHC bond (Fig. 2d), caused the largest separation between TCR and CD3ζ (Fig. 3 e-f), and led to the most complete dissociation of CD3ζ from the inner leaflet of the plasma membrane to maximize the exposure of ITAMs on CD3ζ for subsequent phosphorylation (Fig. 3 i-k). By contrast, the least potent 102S formed the longest bond with the TCR (Fig. 2d), caused the smallest TCR-CD3ζ conformational change (Fig. 3 e-f) and resulted in the weakest phosphorylation of CD3ζ (Fig. 3i-k). Thus, our transmembrane FRET not only directly visualized TCR-CD3ζ conformational changes *in situ*, but also explained the molecular mechanism of signal propagation through CD3ζ phosphorylation for TCR ligand discrimination.

**Fig. 3.**
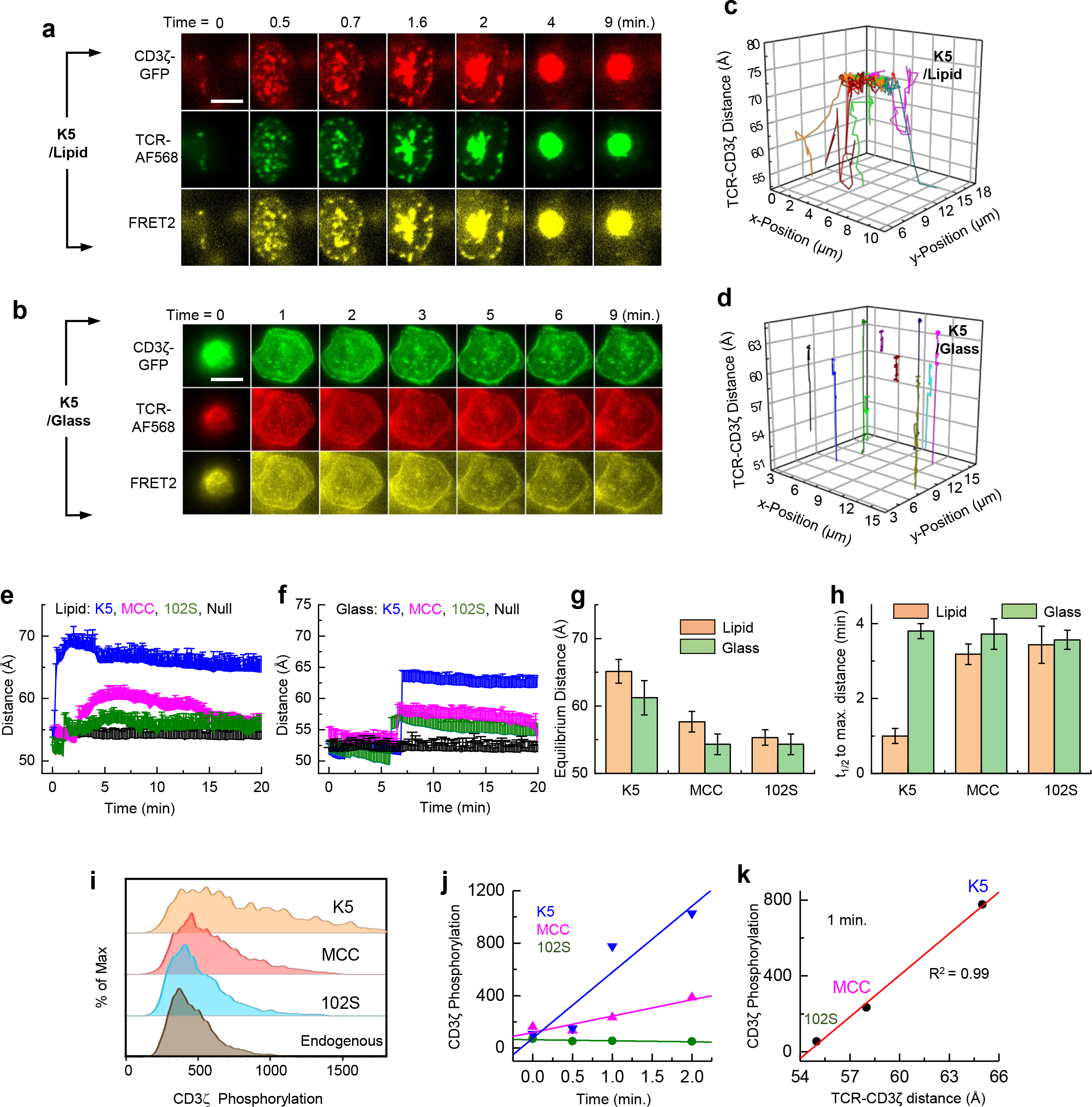
TCR-CD3ζ conformational changes induced by TCR ligation. **a-b**, Representative real-time CD3ζ-GFP (green, donor), TCR-Alexa568 (red, acceptor), and FRET (yellow) signals in a typical transmembrane FRET experiment on lipid bilayer (**a**) or glass surface (**b**) at 37 °C for K5 pMHC. 6-8 independent experiments were repeated for each pMHC on both lipid and glass surface. Scale bar = 5 μm. **c-d**, 3D illustration of the lateral movement (x-y) and intramolecular distance change (z) between TCR and CD3ζ of individual TCR/CD3ζ microclusters shown in Fig. 3a on the lipid bilayer (**c**) and Fig. 3b on the glass surface (**d**). Also see Supplementary Movie 5-7 for microcluster formation, Extended Data Fig. 9–11 for data analysis, Extended Data Fig. 12 for data calibration and Extended Data Fig. 13 for negative controls. **e-f**, Time-dependent TCR-CD3ζ intramolecular distance changes on lipid bilayer (**e**) and glass surface (**f**) upon TCR ligation with K5, MCC, 102S or null pMHC. Each curve showed the average intramolecular distances at consecutive time points from three independent measurements. **g**, The equilibrium intramolecular TCR-CD3ζ distances induced by different pMHCs on lipid bilayer and glass surface. **h**, The half time (t_1/2_) to reach the maximum TCR-CD3ζ distance for different pMHCs on lipid bilayer and glass surface. **i**, Phospho flow showing the phosphorylation of CD3ζ in T cells upon contacting antigen-presenting cells loaded with 102S, MCC or K5 peptides for 1 minute. **j**, The time course of CD3ζ phosphorylation upon stimulation. **k**, Correlation between CD3ζ phosphorylation (1 min stimulation) and TCR-CD3ζ distance.

We then designed a series of experiments to test how TCR-pMHC bond length controls TCR binding, signaling and activation. To test how the TCR-pMHC bond length regulates TCR-pMHC interaction, we performed micropipette adhesion assays to measure the *in situ* two-dimensional (2D) TCR-pMHC binding kinetics and affinities^4^ (Fig. 4a-d, Table-1, Extended Data Fig. 2b, Extended Data Table 1 and Supplymentary Movie 8). As revealed by the co-existence of high binding affinity and short bond length, our data suggested that higher binding affinity drives the formation of more compact TCR-pMHC bond, consistent with the classic bond length theory in biochemistry^21^. To determine how TCR-pMHC bond length controls T-cell signaling, we devised a fluorescent micropipette to measure the real-time T-cell calcium signaling at the single cell level (Fig. 4e-f, Extended Data Fig. 14 and Supplymentary Movie 9). We further adapted the values of half-maximal T-cell proliferation (EC_50_) and 3D half-lives of tetramer binding from reference 9. We then plotted the TCR-pMHC bond lengths and TCR-CD3ζ intramolecular distances against 2D on-rates, 2D affinities, signaling, and proliferation and 3D tetramer half-lives (Extended Data Fig. 15). Strong negative (Fig. 4g top row, solid dots and lines) and positive (Fig. 4g bottom row, open dots and dashed lines) correlations were found between intermolecular TCR-pMHC bond length and intramolecular TCR-CD3ζ distance versus all metrics of TCR binding, signaling and activation, respectively. These measurements provided direct physiological relevance to our studies of TCR-pMHC bond length and TCR-CD3ζ conformational change.

**Fig. 4.**
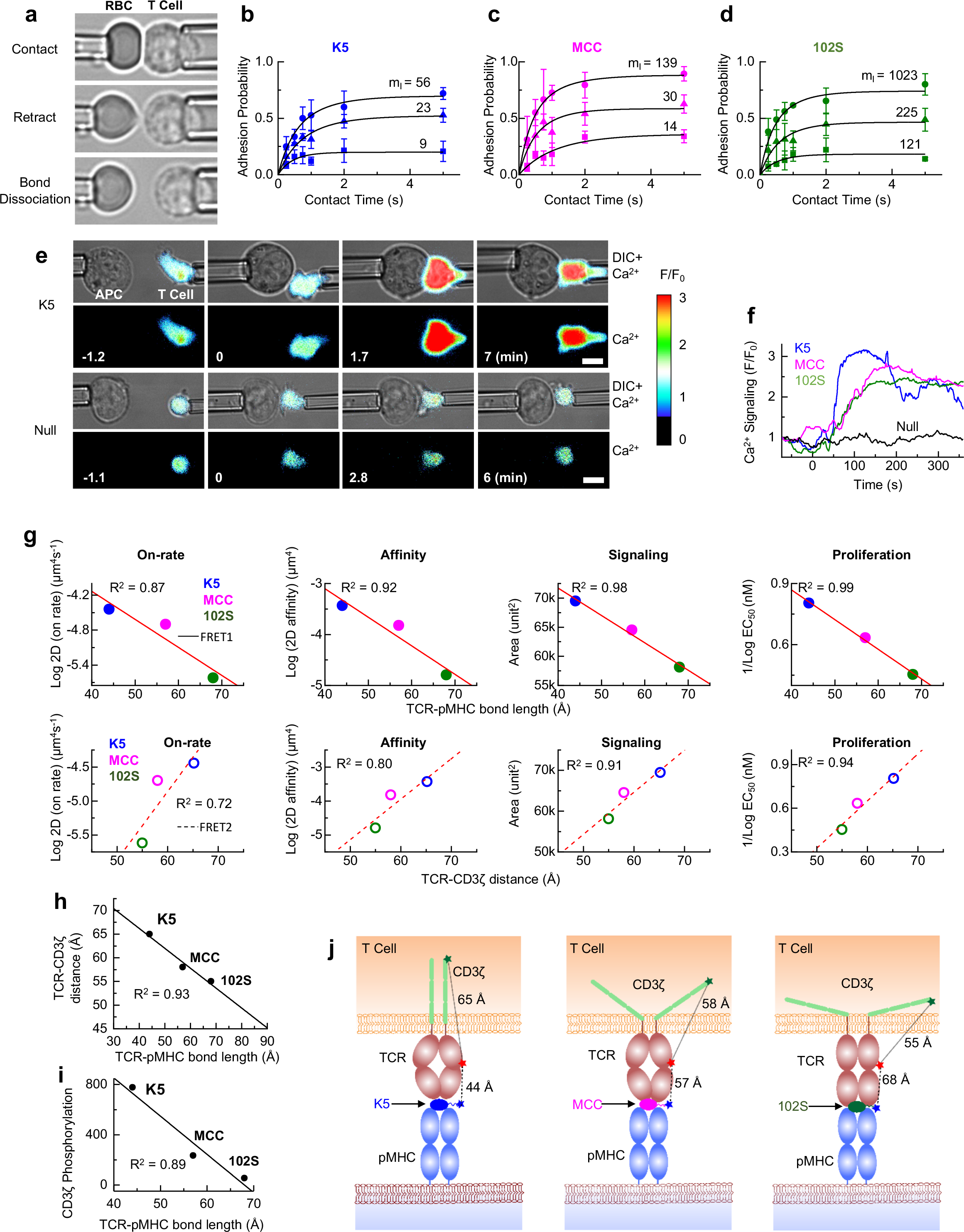
Correlations between TCR conformations and T-cell binding kinetics, signaling and proliferation. **a**, 2D micropipette adhesion frequency assay. A micropipette-aspirated T cell (right) was driven by a piezoelectric translator to make a controlled contact with a RBC coated with pMHC held by another pipette (left). The retraction of T cell to the starting position resulted in elongation of the RBC, enabling visual detection of a TCR-pMHC bond. Also see Extended Data Fig. 1b-c. **b-d**, Adhesion curves for the 5C.C7 TCRs interacting with K5 (**b**), MCC (**c**) and 102S (**d**) pMHCs measured by micropipette at 25 °C at indicated pMHC site densities. Each cell pair was tested fifty times at a given contact duration to estimate an adhesion probability, and 3 cell pairs were tested for each contact duration to calculate a mean adhesion probability. The data (points) were fitted by a probabilistic kinetic model (curves) to determine 2D binding kinetics. Data were summarized in Table 1 and Extended Data Table 1. **e**, Real-time single T-cell calcium signaling measured by fluorescent micropipette. A CH27 cell loaded with K5 (top row) or null peptide (bottom row, control) was precisely controlled to contact a primary 5C.C7 T cell loaded with Fluo-4 calcium indicator at 37 °C. The fluorescence signal was real-time recorded by time-lapse microscopy and the fold-increase of Ca^2+^ signaling (F/F_0_) was shown by pseudo color. Shown were representative Ca^2+^ imaging experiments for K5 and null peptides out of 6-8 independent experiments for each peptide. Also see Extended Data Fig. 14 for peptides MCC and 102S. See Supplementary Movie 9 for more data. **f**, Representative time trajectories of Ca^2+^ signaling stimulated by K5, MCC, 102S and null peptides. Fluorescent intensity values (F) at any given time were divided by the initial fluorescent intensity value at time zero (F_0_) to obtain the fold-increase of Ca^2+^ signaling after cell contact. **g**, Correlations of TCR-pMHC bond length (top row) and TCR-CD3ζ distance (bottom row) with 2D on-rate, 2D affinity, calcium signaling and proliferation for K5, MCC and 102S peptides. EC_50_ data of cell proliferation were adapted from Corse et al. 2010. Data points were fitted with a liner function and the goodness of correlation was indicated by R2 values. **h**, Correlation between TCR-pMHC bond length and TCR-CD3ζ distance. **i**, Correlation between TCR-pMHC bond length and CD3ζ phosphorylation. **j**, Our proposed bond length model for TCR recognition. The TCR-pMHC bond length controls TCR-CD3 distance to regulate the exposure of ITAMs on CD3ζ available for subsequent phosphorylation.

In summary, our data together suggested a “bond length model” in which a TCR deciphers the structural differences between foreign- and self-peptides by forming TCR-pMHC bonds with different lengths (and strengths), which proportionally pull the otherwise sequestered CD3ζ off the inner leaflet of the plasma membrane (Fig. 4h) to expose appropriate degrees of ITAMs for subsequent phosphorylation (Fig. 4i) in a precisely controlled manner. Mechanical forces^4,22–24^ and calcium feedback regulation^25^ are the most likely mechanisms of CD3ζ sequestration off the inner leaflet. Our “bond length model” (Fig. 4j) is compatible with existing TCR triggering models^6,26^, can well explain previous studies of CD3 conformational changes^27–30^ and suggests that T cells use simple, reliable and efficient machinery to faithfully translate extracellular TCR-pMHC binding to proper intracellular signaling for optimal T-cell triggering. Our results revealed the dynamic process of how the recognition signal is initiated, controlled and transmitted by directly linking intermolecular TCR-pMHC bond length and intramolecular TCR-CD3ζ distance to T-cell surface binding, intracellular signaling and functional response, thus explaining the molecular mechanism of TCR recognition at the very first step.

**Table 1.** 2D kinetic parameters.

| Peptide | Sequence | T cell activation | $A_c K_a (\mu\text{m}^4)$ | $k_r (\text{s}^{-1})$ | $A_c K_{\text{on}} (\mu\text{m}^4 \text{s}^{-1})$ |
| --- | --- | --- | --- | --- | --- |
| K5 | ANERADLIA<br>YFKAATKF | Super Agonist | $3.7 \pm 0.6 \times 10^{-4}$ | $1.0 \pm 0.3$ | $3.6 \pm 0.5 \times 10^{-5}$ |
| MCC | ANERADLIA<br>YLKQATK | Agonist | $1.5 \pm 0.7 \times 10^{-4}$ | $1.3 \pm 0.7$ | $2.0 \pm 1.1 \times 10^{-5}$ |
| 102S | ANERADLIA<br>YLKQASK | Weak Agonist | $1.6 \pm 0.7 \times 10^{-5}$ | $1.5 \pm 0.3$ | $2.4 \pm 0.8 \times 10^{-6}$ |

## Acknowledgements

We thank Mark M. Davis at Stanford for providing the constructs of anti-TCR single-chain variable fragments, Michael Birnbaum at MIT for providing IE^k^ plasmids, Xiaolei Su and Marcus Taylor at UCSF for advice in the preparation of glass supported lipid bilayer and the NIH Tetramer Core Facility for providing pMHC monomers. This work was supported by NIH grants R00AI106941 and R21AI120010, NSF CAREER Award 1653782, and Chicago Biomedical Consortium Catalyst Award (to J.H.) and postdoctoral grant PDR-092 (to D.K.S.) with support from the Searle Funds at The Chicago Community Trust.

## Author Contributions

J.H. conceived, directed and supervised the project. J.H. and D.K.S. designed experiments. D.K.S. custom built the fluorescence microscopy system, performed imaging experiments, and analyzed data unless indicated otherwise. W.F. and D.K.S. custom built the micropipette system. W.F. performed all 2D binding kinetics assays and analyzed the data. W.F. and D.K.S. performed Ca^2+^ imaging experiments and D.K.S. analyzed data. Reagents and experimental systems were designed and tested by D.K.S. unless indicated otherwise. S.R. and E.A. generated IE^k^ proteins. H.L. and J.Q. performed PMF analysis. G.C. analyzed smFRET data. Y.H. and H.S. prepared the CD3ζ-GFP construct and infected primary T cells. C.C. performed phospo flow cytometry. D.K.S. and P.L. cultured T cells. D.K.S. and J.H. wrote the manuscript with input from all authors.

## Additional information

### Extended data

This section contains 15 Extended Data Figures and captions.

### Supplymentary Information

This section contains Supplymentary Movies 1-9 and captions.

## METHODS

### Mice

The Institutional Animal Care and Use Committee of the University of Chicago approved the animal protocols used in this study. 5C.C7 TCR-transgenic RAG2 knockout mice in B10.A background were a generous gift from NIAID.

### Cells

5C.C7 T cell blasts were obtained by stimulating lymph node and spleen cells from transgenic mice with 10 μM MCC peptide (amino acids 88–103, ANERADLIAYLKQATK). T cell blasts were maintained in complete medium (RPMI 1640 medium, 10% fetal calf serum, 2 mM L-glutamine, 50 μM β-mercaptoethanol, and penicillin-streptomycin). T-cell blasts were used on days 6–9 for imaging and micropipette experiments. Live T cells were separated from dead cells by Ficoll-Paque density gradient media (GE). The B-cell lymphoma cell line CH27 was used as APCs. APCs were maintained in the same medium as T cells^2,3^. Human RBCs were isolated from whole blood of healthy donors from the Hospital of the University of Chicago according to protocols approved by the Institutional Review Board of the University of Chicago.

### Reagents

1-palmitoyl-2-oleoyl-*sn*-glycero-3-phosphocholine (POPC), 1,2-dioleoyl-*sn*-glycero-3-[(N-(5-amino-1-carboxypentyl) iminodiacetic acid) succinyl] nickel salt (DGS-NTA-Ni^2+^) and 1,2-dioleoyl-*sn*-glycero-3-phosphoethanolamine-N [methoxy (polyethyleneglycol)-5000] ammonium salt (PEG5000 PE) were purchased from Avanti Polar Lipids. PBS, BSA, FBS, and Alexa Fluor 568 C5 Maleimide were purchased from Thermo Fischer Scientific. PEG-NTA-Ni^2+^ coated cover glasses were purchased from MicroSurfaces, Inc. Nunc Lab-Tek 8-well chambered cover glasses were purchased from Thermo Fisher Scientific. His-tagged B7 was made as previously reported^3^. His-tagged ICAM-1 was purchased from Sino Biological. Cy3 and Cy5 maleimide mono-reactive dyes were purchased from GE Life Sciences. Alexa-568, Fura-4 and di-methyl sulfoxide (DMSO) were purchased from Thermos Scientific. LB-media were from Fisher Scientific.

### pMHCs

For FRET experiments between TCR and pMHC on the cell surface (FRET1), we generated peptide exchangeable CLIP-IE^k31^. The α and β chains of CLIP-IE^K^ were cloned in pAC vectors with gp67 signal sequence (BD biosciences), acidic or basic zipper sequences and a 6-histidine tag at the C terminus. Primary baculoviruses were prepared for each chain by co-transfecting the construct with linearized Baculovirus DNA (BestBac 2.0, Expression Systems) into the Sf9 cells using cellfectin reagent (Thermo Fischer Scientific), cells were washed and incubated at 27 °C for a week. Primary viruses were harvested by centrifugation and collecting the supernatant. Baculoviruses were amplified for higher titer by infecting Sf9 cells for another week. Hi5 cells were co-infected with baculovirus for both chains and supernatant was harvested after 65 hours of infection. The pH of supernatant was adjusted to pH 6.9 with HEPES buffer saline (10 mM HEPES pH 7.2, 150 mM NaCl, 0.02% NaN3) and 20 mM imidazole pH 7.2, 5 mM MgCl_2_, 0.5 mM NiCl_2_. Ni-NTA agarose (Qiagen) was added to the supernatant and stirred overnight at 4 °C. Supernatant was filtered, Ni-NTA agarose was collected and CLIP-IE^K^ was eluted using 200 mM imidazole, pH 7.2 in HBS. Protein fractions were analyzed in SDS-PAGE. CLIP-IE^K^ was purified using Superdex 200 size exclusion column chromatography and Mono-Q anion exchange chromatography (GE healthcare). Purified fractions were used for peptide loading. According to previous publication^3^, peptides with fluorescent maleimide dye (Cy3) at the C-terminus, including K5(C)-ANERADLIAYFKAATKdFGGdSdC, MCC(C)-ANERADLIAYLKQATKGGdSdC, T102S(C)-ANERADLIAYLKQASKGGdSdC, null(C)-ANERAELIAYLTQAAKGGGdSdC were synthesized, labeled with Cy3, purified by HPLC and confirmed by mass spectroscopy by ELIM BIOPHARM company, CA. Peptides of interest were added to the purified CLIP-IE^k^ protein at 100-fold molar excess for the peptide exchange reaction. The thrombin (1 U /100 μg of IE^k^) was added and incubated at 37 °C for one hour. The pH of the solution was lowered by adding MES buffer, pH 6.2 to a final concentration of 30 mM, and IE^K^ was again incubated at 37 °C overnight. The pH of protein solution was adjusted with 40 mM HEPES, pH 7.2. Extra peptides and denatured proteins were removed by centrifugation (16,000×g for 30 min at 4 °C) and desalting twice with Zeba Spin Desalting Columns (Thermo Fisher).

For FRET experiments between TCR and CD3ζ across the cell membrane (FRET2) and 2D micropipette adhesion assays, we used IE^k^ pMHC monomers generated by the NIH Tetramer Core Facility. Biotinylated, monomeric IE^k^ were covalently complexed with ANERADLIAYFKAATKF (K5), ANERADLIAYLKQATK (MCC), ANERADLIAYLKQASK (102S) and PVSKMRMATPLLMQA (human CLIP 87-101) peptides. IE^k^ monomers were aliquoted and stored at −80 °C for experiments.

### Production and Labeling of Single Chain Variable Fragments (scFvs)

Plasmid constructs of two mutants of anti-TCR scFvs J1 and J3 were a generous gift from Mark M. Davis at Stanford University^3^. J1 was used for cell surface FRET1 and J3 was used for transmembrane FRET2 experiments, respectively. To generate J1 and J3 proteins, BL21 bacteria were transfected with cDNA and cultured in large scale (2 L each) in the presence of Isopropyl β-D-1-thiogalactopyranoside. Bacteria were spun down, and cell pellets were resuspended with B-PER (Bacterial Protein Extraction Reagent, Thermo Fisher), lysozyme, and DNase, followed by washing with inclusion wash buffer (100 mM NaCl, 50 mM Tris Base, 0.05% (volume) Triton X-100)^32^. Refolding and purification of recombinant scFvs were performed using a modified method based on previous publications^33,34^. In brief, scFvs were unfolded in the presence of 10 mM β-mercaptoethanol for 2 hours at 25 °C (in 100 mL of 100 mM Tris-HCl buffer with pH = 8.0, 6 M GuHCl and 200 mM NaCl), so that it could be refolded in correct conformations by step-wise dialysis methods without causing protein oxidation. To remove the reducing agent, denatured recombinant scFvs were dialyzed against 1 L of 100 mM Tris-HCl, pH 8.0 buffer with 6 M GuHCl and 200 mM NaCl for 15 hours at 4 °C with gentle stirring. After that, stepwise dialyses were performed in the same Tris-HCl buffer containing decreasing GuHCl concentrations (4 M, 2 M, 1 M, 0.5 M and 0 M) for 15 hours each step at 4 °C with gentle stirring. During 1 M and 0.5 M dialysis steps, 400 mM L-arginine (Sigma-Aldrich) and 375 μM of oxidized glutathione (Sigma-Aldrich) were added. Final dialysis was performed in buffer without GuHCl for 18 hours at 4 °C with gentle stirring. Protein was concentrated and stored at 4 °C before long-term storage at −20°C in the presence of 50% Glycerol. Purified scFVs were labeled with Cy5-malimide in the presence of 50 μM of tris-(2-carboxyethyl) phosphine hydrochloride (TCEP) for two hours at room temperature followed by 12 hours gentle mixing at 4 °C. Then the labeled scFVs were purified by resin spin column. J1 and J3 were labeled and purified freshly before each experiment. The binding specificity of scFvs for 5C.C7 TCRs was confirmed by Flow cytometry before imaging applications.

### CD3ζ-GFP Transduction

Primary 5C.C7 T cells were retrovirally transduced with CD3ζ-GFP according to a previously published method^35^. Ecotropic platinum-E retroviral packaging cells were transiently transfected with MIG-CD3ζ-GFP vector by calcium phosphate precipitation. Viral supernatant was harvested twice at 48 hours and 72 hours post transfection, filtered by 0.2 μm cellulose acetate membrane and used for following experiments. Splenocytes isolated from 5C.C7 mice were cultured in RPMI supplemented with 10% FCS, 2 mM glutamine, 50 μM 2-mercaptoethanol, 1 mM HEPES, 1 mM sodium pyruvate, 1Χ glutamine and non-essential amino acids (Thermo Fisher), 100 U/mL penicillin, 100 μg/mL streptomycin, and 50 μg/mL gentamycin, and stimulated with pre-coated 1.5 μg/mL anti-CD3ɛ Ab (Clone 145-2C11, University of Chicago Monoclonal Antibody Facility) and 0.5 μg/mL anti-CD28 Ab (Clone 37.51, Biolegend) in the presence of 40 U/mL recombinant human IL-2 (Peprotech). After 24 hours of cell activation, 2 mL viral supernatant were spun down in a well of a Retronectin (Clontech) pre-coated (12.5 μg/mL in PBS, 4C overnight) 6-well plate for 90 min at 3,000 g, and then the stimulated cells were transferred to the plate and spun down in the viral supernatant supplemented with 4 μg/mL protamine sulfate at 800 g for 90 min. The transduction rate of CD3ζ-GFP was determined by GFP fluorescence at least 16 hours after transduction.

### Lipid Bilayer

The glass-supported lipid bilayer preparation method was developed based on previous publications^15,36^. Lipid layer was made by mixing POPC (90%), DGS-NTA-Ni^2+^ (9.9%) and PEG500PE (0.1%) in chloroform in clean glass vials. Chloroform was dried by blowing 0.22 μm filtered N_2_, and then vials containing lipid layer were kept in vacuum for two hours to dry completely. The lipid layer was then resuspended in filtered PBS (pH 7.4) buffer (Clontech) at a concentration of 4 μmol/mL. To decrease the size of the multilamellar lipid vesicles to unilamellar vesicles, the cloudy vesicle solution was repeatedly frozen in liquid nitrogen and thawed in 37 °C water bath 30 times until the solution become clear. The unilamellar vesicle solution was stored in −80 °C for future experiments. Before each experiment, a tube was centrifuged at 33,000×g for 45 minutes at 4 °C. The supernatant was incubated for 90 minutes on the 8-well Lab-Tek chamber cover glass that was prewashed thoroughly by 5N NaOH twice at 50 °C and followed by PBS twice at 37 °C. The lipid vesicles fused on the glass surface and formed the glass supported lipid bilayer. The lipid bilayer was washed three times thoroughly with PBS to remove excess lipid. Then a mixture of his-tagged, fluorescently-labeled pMHC and his-tagged, non-fluorescent ICAM-1 and B7 was added to the lipid bilayer and incubated for 1 hour. After 30 min, unbound proteins were washed off by PBS for three times. The bound protein was incubated for another 30 min at 37 °C, and weakly bound proteins were washed off with PBS for an additional three times. The protein-bound lipid was incubated with 1% BSA for 20 minutes to minimize background fluorescence during microscopy experiments. Excess BSA was washed three times with PBS. Microscopy experiments were performed using imaging buffer: PBS pH 7.4, 137 mM NaCl, 5 mM KCl, 1 mM CaCl_2_, 2 mM MgCl_2_, 0.7 mM Na_2_HPO_4_, 6 mM D-glucose and 1% BSA.

The fluidity and integrity of lipid bilayer were tested by fluorescence recovery after photobleaching (FRAP) experiments with 32 nM Cy3-labeled pMHC reconstituted on supported lipid bilayer (Extended Data Fig. 3 and Supplymentary Movie 2). Photobleaching was performed by a high power (60 mW) 532-nm CW-laser for 5 second exposure at the center of the imaging area, and the fluorescence recovery of lipid bilayer was imaged by a 470 ± 10 nm LED light with 10 seconds interval. The power and duration of laser and LED light excitation were controlled by analog modulation. The diffusion coefficient (*D*) was determined by labeling the lipid bilayer with 1 nM of Cy3-labeled pMHC. The experimentally determined diffusion coefficient *D*_e_ was verified by a small-scale simulation *D*_s_ which showed excellent resemblance (Extended Data Fig. 4 and Supplymentary Movie 1). pMHC diffusion constant was determined by TrackArt programming in MatLab, consistent with the previously reported values^37^.

### Microscopy

All imaging experiments were performed using our custom-built total internal reflection fluorescence (TIRF) and epi-fluorescence microscope setup, which was based on a Nikon-Ti-E inverted microscope attached with an Optosplit-III (CAIRN Research) and an Andor iXon 888 EMCCD camera (1024×1024 pixel) (Extended Data Fig. 2a). Individual CW laser lines of 405-nm, 488-nm, 532-nm and 647-nm (Cobolt) were aligned to achromatic fiber port (APC type, 400-700 nm Thorlabs, Inc) and then passed through an achromatic polarization maintaining single-mode fiber to a motorized Nikon TIRF illuminator. The lasers were reflected by a custom-built quad-band dichroic mirror (Chroma, ZT405-488-532-640rcp) to the sample through a 1.42 NA 100× TIRF objective. A 7-color solid state LED light source with band-pass filters was also attached with a liquid light guide to the upper filter cube wheel in the Nikon microscope. The fluorescence from donor and acceptor passed through quad band laser clean-up filter (ZET405-488-532-647m) and then passed through Optosplit III to separate the emission fluorescence of FRET donor and acceptor. In optosplit cube, we used different filter sets for FRET1 and FRET2 experiments. For FRET1 (Cy3-Cy5), we used T640lpxr-UF2 (Chroma) as dichroic and ET585/65 for Cy3 channel (Chroma) and ET655lp for Cy5 channel (Chroma) for individual fluorescence signal. For FRET2 (GFP-Alexa568), we used dichroic T560lpxr-UF2 and two band pass filters Chroma ET510/20m for GFP and Chroma ET595/50m for Alexa-568. For single-molecule imaging, we applied hardware sequencing to get the highest frame rate (5-10 ms exposure, 90-210 frame/second) using high EM gain. Either hardware sequencing (by camera) or software (Micromanager) sequencing was applied with analog modulation to synchronize the image acquisition by EMCCD camera that triggers each laser and individual LED source. The donor and acceptor signals channels were physically separated out by the beam splitter in Optosplit-III and both signals were imaged simultaneously on the same image frame. Hardware stage control and images were acquired by Micromanager^38^.

### 2D Fluorescent Micropipette

The micropipette apparatuses were centered around a Leica inverted microscope placed on an anti-vibration table (Newport) equipped with manometer systems to apply suction pressures through glass pipettes (Extended Data Fig. 2b). Two opposing pipettes are attached to two identical piezoelectric micromanipulators (Sensapex) to control contacts between a T cell and a pMHC coated RBC or a CH27 APC. In the micropipette apparatus, one of the pipettes was also attached to a PI piezo actuator to allow computer-programmed fine movements for the repeated adhesion test cycles. The cell chamber to the desired size was prepared by cutting cover glass. Temperature of the cell chamber (37 °C) was controlled by an objective heater (Bioptechs). To avoid medium evaporation for heating, the chamber was sealed by mineral oil (Sigma). The real-time images were acquired by an Andor iXon 888 EMCCD camera through 100× objective and Micromanager software. For real-time calcium imaging, the sample was illuminated by sequentially triggered light exposure of 470 ± 20 nm blue light (Spectra X, Lumencor) and white LED light. Triggering of light channels and data acquisition were performed in analog modulation using Micromanager ^38^. For 2D kinetic measurements, only continuous white LED light was used to detect the adhesion between a T cell and a pMHC coated RBC.

### 2D Micropipette Kinetic Assay

2D micropipette adhesion experiments^4^ were performed using T cell blasts^2^ and pMHC-coated RBCs. Monomeric pMHCs were coated onto RBCs by biotin-streptavidin coupling. RBCs isolated from the whole blood were biotinylated using different concentrations of biotin-X-NHS according to the manufacturer’s instructions. 10 μl RBCs solution (~10 × 10^6^) with 2 mg/mL streptavidin solution (10 μl) were mixed for 30 min at 4 °C and then incubated with 20 μg/mL biotinylated pMHC monomers for 30 min at 4 °C. After each step, cells were washed three times.

To determine the surface receptor and ligand densities, T cells were incubated with a PE-conjugated anti-mouse TCR β chain antibody clone H57-597 (BD) at 10 μg/mL in 200 μl of FACS buffer (RPMI 1640, 5 mM EDTA, 1% BSA and 0.02% sodium azide) at 4 °C for 30 min, and pMHC-coated RBCs were stained with a PE-conjugated anti-IE^k^ clone 14.4.4s (BD). T cells and RBCs were analyzed by a BD LSRFortessa flow cytometer. The fluorescence intensities were compared to standard calibration beads (BD Quantibrite PE Beads, BD) to determine the total number of molecules per cell, which were divided by the cell or bead surface area to obtain site densities. The apparent surface areas of T cells blasts (523 μm^2^) and RBC surface area (140 μm^2^) were calculated as the areas of smooth spheres from their radii measured microscopically.

T cells were incubated at 37°C in saturating conditions (10 μg/mL) of purified anti-mouse CD4 (clone GK 1.5, BD) prior to addition to chamber. The RBC was driven in and out of contact with T cell with controlled contact time (0.25, 0.5, 0.75, 1, 2, and 5 s) and area by computer programming. Adhesion events were detected by observing RBC elongation upon cell separation. The contact-retraction cycle was repeated fifty times for a given contact time. Specific adhesion probability (*P*_*a*_) at each contact time point was calculated by subtracting the nonspecific adhesion frequency (*P*_*nonspecific*_). Following equations were used to analyze the data.

*P*_a_ versus contact time *t* were fitted using a probabilistic model (Equation 1)^39^:

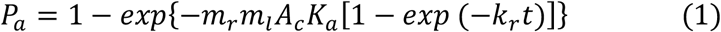

Where *K*_a_ and *k*_r_ are the 2D binding affinity and off-rate, *m*_r_ and *m*_1_ are the respective TCR and pMHC densities that were measured by flow cytometry, and *A*_c_ is the contact area. The curve-fitting generates two parameters, the effective 2D affinity *A*_c_*K*_a_ and the 2D off-rate *k*_r_. Its product with the off-rate is the effective 2D on-rate:

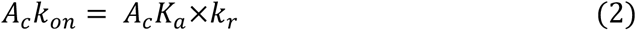

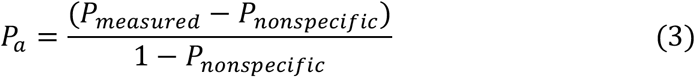

where *P*_nonspecifc_ and *P*_measure_ are the nonspecific adhesion fraction and the total measured adhesion, respectively.

### FRET Analysis

FRET is a nonradiative process and originated by dipole-dipole interaction between the electronic states of donor and acceptor^8,40^ The energy transfer occurs only when the oscillations of an optically-induced electronic coherence of the donor are resonant with the electronic energy gap of the acceptor. Efficiency of energy transfer (*E*_*FRET*_) is sensitive to the inter-distance between the donor and acceptor, which is typically in the range of 10-100 Å. The energy transfer efficiency (*E*_*FRET*_) is given by^8,40^:

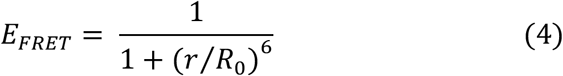

Where *r* is inter-distance between donor (*D*) and acceptor (*A*), and *R*_*0*_ is Förster’s distance. FRET is very sensitive to the distance between the acceptor and the donor, which may change due to conformation dynamics or other factors. The *E*_*FRET*_ between *D* and *A* was calculated by a ratiometric method to reveal the conformational dynamics using the following equation^8,40^:

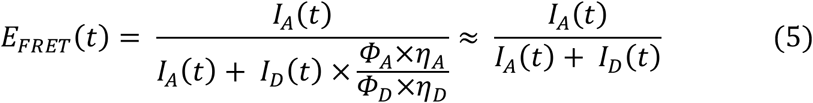

Where *I* is fluorescence intensity, *Φ* is emission quantum yield, *η* is photon detection efficiency. Here the correction factor is ~ 1 in our experimental condition.

To obtain the distribution of the FRET efficiencies, *E*_*FRET*_, and the corresponding distance between a FRET donor and a FRET acceptor, we tracked and measured the fluorescence intensities of the donor and the acceptor. For each experiment, image registration was first performed using MATLAB on the images of the corresponding frame from Donor Channel and Acceptor Channel. The donor and acceptor molecules were identified and tracked using TrackMate in ImageJ until the end of each FRET trajectory. The fluorescence intensities of a donor and an acceptor were measured and background corrected frame-by-frame. Equations 4 and 5 were applied to calculate *E*_*FRET*_ and corresponding distance for both FRET1 (Cy3 and Cy5) and FRET2 (GFP and Alexa568) experiments. By tracking the trajectories of many individual FRET pairs, we obtained the values and distribution of *E*_*FRET*_ and corresponding distance.

### Potential-of-Mean-Force (PMF) Analysis

One way to quantify the binding strength is to examine the potential-of-mean-force (PMF) for the fluctuation of donor-receptor distance *R*. The PMF in this context is given by^12,41^:

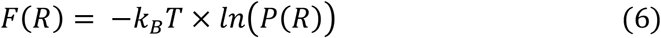

where *P*(*R*) is the histogram of the distance that has been averaged over the steady-state signals collected over 8,000 independent TCR-pMHC binding events. The PMF measures the free energy cost of variation in distance *R*. It is minimized at equilibrium. Its curvature governs the size of fluctuation. Shallower potential implies wilder fluctuation and weak binding.

### Microclusters Tracking Analysis

In FRET2 analysis, we developed a method to track individual TCR and CD3ζ micro-clusters in three dimensions (x, y and z) using TrackMate plugin in Fiji^42^. The track of each individual donor and acceptor cluster gave the lateral movement (x-y axis) as well as FRET2 efficiency (TCR-CD3 distance, z axis) as microclusters moving towards the center and forming the immunological synapse.

### Measurement of CD3ζ phosphorylation

We used phospho flow cytometry to measure the phosphorylation state of CD3ζ at the single cell level^43,44^. CH27 cells were preincubated with 10 μM peptides in complete medium for 3 hours at 37 °C. Plain CH27 cells were used as a negative control. Peptide loaded CH27 cells were washed three times^2^. 5C.C7 T cells were rested in serum free RPMI medium at 37 °C for 3 hours to reduce the phosphorylation background^15^. 50,000 peptide-loaded CH27 cells and 50,000 rested 5C.C7 T cells were precooled and mixed in a tube on ice. The tube was centrifuged at 300× g for 1 min at 4 °C to initiate cell-cell contact, and immediately transferred to a 37 °C water bath for initiating T-cell stimulation. The stimulation was terminated at indicated time points with 4% PFA fixation. After 10 minutes fixation at room temperature, cells were washed twice with ice-cold PBS containing 2% BSA, and then resuspended in 80% methanol and incubated for 30 min at −20 ̊C. After washing twice with ice-cold PBS, Alexa488 labeled anti-pY142-CD3ζ antibody (BD) was added in a final volume of 100 μl of ice-cold PBS and incubated at 4 ̊C for 45 min. Cells were washed three time with ice-cold PBS containing 2% BSA and analyzed by Fortessa flow cytometry. Flow data were further processed with FlowJo software.

### Ca^2+^ Imaging

For Ca^2+^ flux experiments, T cells (~10^6^) were incubated with 5 μM of fluorescent dye Fluo-4 AM (Molecular Probe) for 30 min in complete RPMI-1640 medium. All the Fluo-4 loading and imaging experiments were performed in the presence of 2.5 mM probenecid. T cells were washed twice with minimal imaging media (MIM, colorless RPMI with 5% FCS and 10 mM HEPES), and then transferred to MIM for 10 min at 37 °C before data collection^45^. For imaging, a LEITZ DMIRB Leica Microscope was used equipped with a 100× objective and an iXON Ultra 888 EMCCD Camera. Calcium flux imaging acquisition was made with Micromanager software. For T cell-APC conjugate experiments, CH27 B cells (10^6^) were pulsed with 4 μM of each peptide for 4 hours at 37 °C, and then washed with MIM. T cells (2 μL) and CH27 B cells (2 μL) were added into MIM (300 μL) in the cell chamber. Seal the chamber using mineral oil from both sides to avoid MIM evaporation. Signals from Fluo-4 were collected at intervals of 100 ms for up to ~20 min and post-processed by Fiji.

## Supplymentary Information

### Supplementary Movie Legends

**Supplementary Movie 1 | Single pMHC molecules diffusion on glass supported lipid bilayer** |His_12_-K5(Cy3)-IE^k^ molecules were attached to Ni^2+^-functionalized glass supported lipid bilayer at 32 nM. IE^k^ pMHC diffusion was imaged by TIRF microscopy with 50 ms exposure at 37 °C. Shown is a representative video out of eight independent experiments. Tracking paths of single IE^k^ pMHC diffusion on lipid bilayer were shown on right side. The video was set to play at 50 frames/second with a field of view of 30 × 27 μm. Scale bar: 5 μm.

**Supplementary Movie 2 | Fluorescence recovery after photobleaching (FRAP)** | His_12_-K5(Cy3)-IE^k^ molecules were loaded to Ni^2+^-functionalized glass supported lipid bilayer. Time-lapse microscopy experiment showed the entire process of pre-bleaching, bleaching, and post-bleaching recovery. The diffusion of lipid bound protein recovered 60-80% of the intensity at the photo-bleached area. FRAP were imaged by epi-fluorescence microscopy with a 200 ms exposure and 10 second interval for 6.8 min at 37 °C. Shown is a representative video out of seven independent experiments. The video was set to play at 1 frame/second with a field of view of 37 × 38 μm. Scale bar: 5 μm.

**Supplementary Movie 3 | Synapse formation** | His_12_-K5(Cy3)-IE^k^, His_12_-B7 and His_12_-ICAM molecules were attached to Ni^2+^-functionalized glass supported lipid bilayer. Time series of synapse formation were recorded by epi-fluorescence microscopy with an interval time of 40 ms at 37 °C. Fluorescence (right) and fluorescence overlaid with DIC (left) channels were shown side by side. Shown is a representative video out of 3 independent experiments. The video was set to play at 15 frames/second with a field of view of 37 × 38 μm. Scale bar: 2 μm.

**Supplementary Movie 4 | Single-molecule FRET** | His_12_-K5(Cy3)-IE^k^, His_12_-B7 and His_12_-ICAM were attached to Ni^2+^-functionalized glass supported lipid bilayer. TCR was labeled with scFv J1-Cy5. Time series of FRET signals were imaged by TIRF microscopy with an exposure time of 10 ms at 37 °C. Donor and acceptor fluorescence signals were simultaneously recorded by a same EMCCD. Shown is a representative video out of 10-15 independent experiments. The video was set to play at 15 frames/second with a field of view of 37 × 38 μm. Scale bar: 2 μm.

**Supplementary Movie 5 | CD3ζ-GFP microcluster formation on lipid bilayer** | His_12_-K5-IE^k^, His_12_-B7 and His_12_-ICAM were attached to Ni^2+^-functionalized glass supported lipid bilayer. Time series of CD3ζ-GFP microcluster formation and diffusion were imaged by epi-fluorescence microscopy with a 40 ms interval at 37 °C. Shown is a representative video out of 6-8 independent experiments. The video was set to play at 11 frames/second with a field of view of 12 × 14 μm. Scale bar: 3 μm.

**Supplementary Movie 6 | CD3ζ-GFP microcluster formation on glass surface** | His_12_-K5-IE^k^, His_12_-B7 and His_12_-ICAM were attached to Ni^2+^-functionalized PEG coated glass. Time series of CD3ζ-GFP microcluster formation and diffusion were imaged by epi-fluorescence microscopy with a 40 ms interval at 37 °C. Shown is a representative video out of 6-8 independent experiments. The video was set to play at 11 frames/second with a field of view of 18 × 22 μm. Scale bar: 3 μm.

**Supplementary Movie 7 | TCR microcluster diffusion to the center of synapse on lipid bilayer** | His_12_-K5-IE^k^, His_12_-B7 and His_12_-ICAM were attached to Ni^2+^-functionalized glass supported lipid bilayer. TCRs were labeled with Alexa568-scFv J3. TCR microcluster formation and diffusion to the center of synapse was imaged by epi-fluorescence microscopy with an interval time of 255 ms at 37 °C. Shown is a representative video out of 6-8 independent experiments. The video was set to play at 51 frames/second with a field of view of 63 × 63 μm. Scale bar: 10 μm.

**Supplementary Movie 8 | Measuring *in situ* kinetics and affinity of TCR-pMHC interactions by 2D micropipette adhesion assay** | Biotin pMHCs were coated onto red blood cell (RBC) surface at an optimal ligand density. A 5C.C7 transgenic T cell and a pMHC coated RBC were brought in and out for 50 times to observe the adhesion frequency at a pre-defined contact duration at 25 °C. The RBC acts as an adhesion sensor by stretching its membrane in response to the force (if adhesion is present) when the T cell is retracted, enabling visualization of adhesion. Scale bar: 10 μm.

**Supplementary Movie 9 | Single-cell Ca^2+^ Imaging** | Peptides (K5, MCC, 102S and Null) were loaded onto CH27 cells (APC) surface at 4 μmol concentration. T cells were loaded with Fluo-4 Ca^2+^ sensor dye for 30 mins at 37 °C. The APC and the T cell were brought into contact, and the real-time Ca^2+^ flux was imaged by epifluorescence microscopy at an interval of 87 ms at 37 °C. Shown is a representative video out of 3-5 independent experiments for each pMHC (K5, MCC, 102S and null). The video was set to play at 15 frames/second with a field of view of 61 × 104 μm for each pMHC. Scale bar: 5 μm.

**Extended Data Fig. 1.**
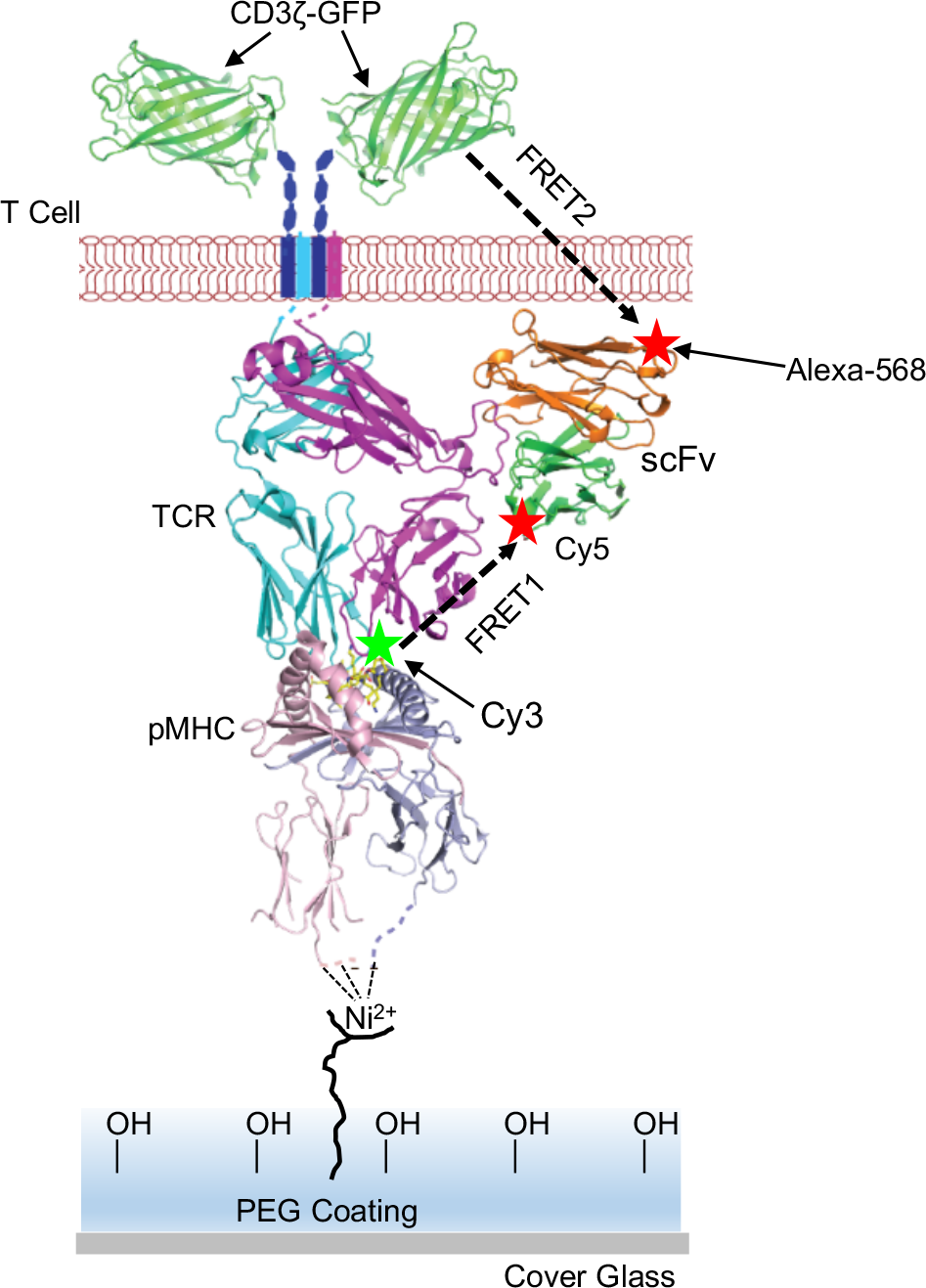
FRET on glass surface. His-tagged pMHCs and accessory molecules ICAM-1 and B7.1 were anchored on the PEG-Ni^2+^ glass surface to perform cell surface FRET (FRET1) and transmembrane FRET (FRET2) experiments. Also see Fig. 1a when pMHCs and accessary molecules were anchored on lipid bilayer.

**Extended Data Fig. 2.**
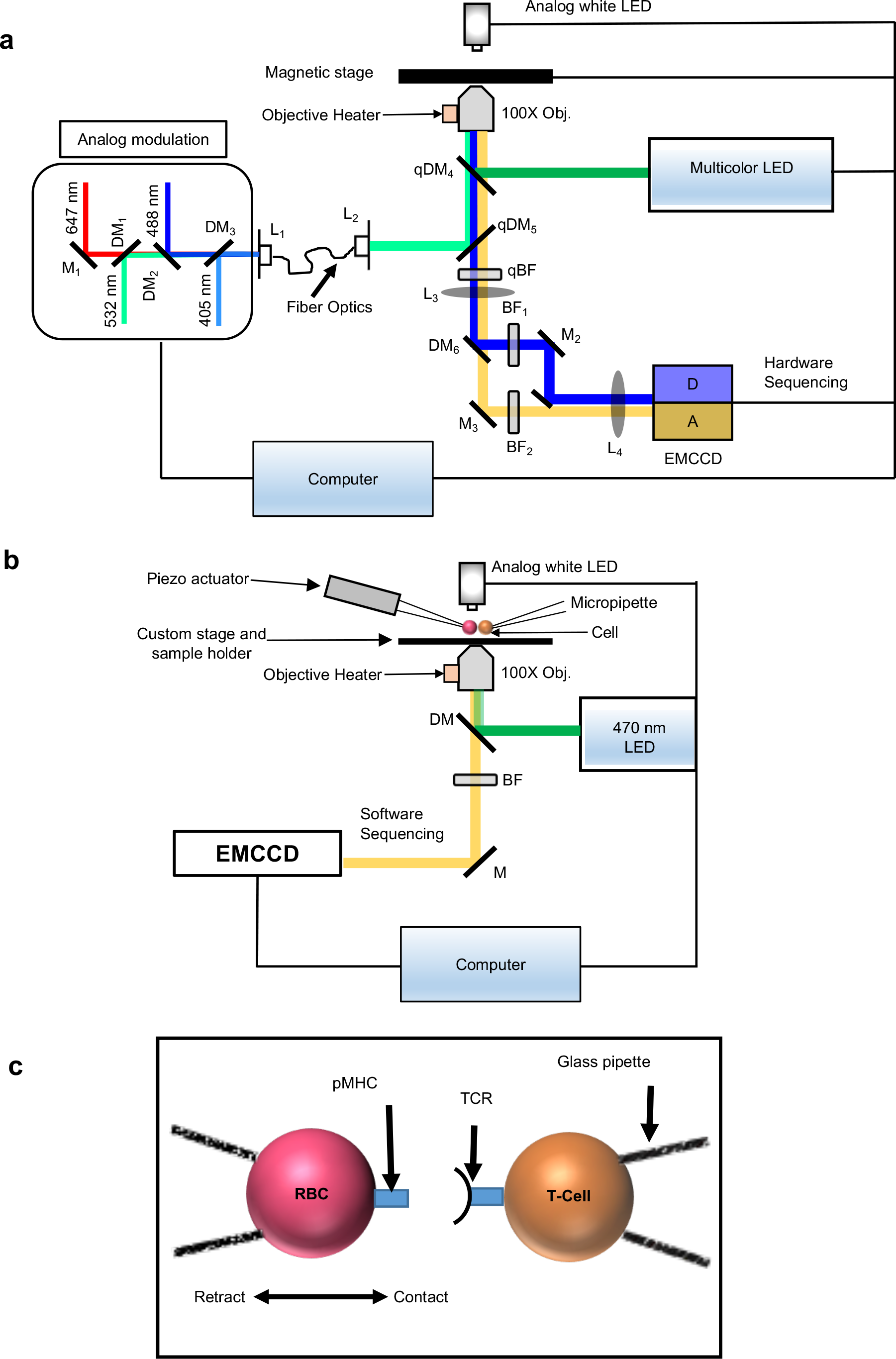
Schematics of fluorescence microscopes. **a**, The diagram illustration of a dual-function TIRF and epi-fluorescence microscope. DM: Dichroic mirror; qDM: quad-band dichroic mirror; L: Lens; qBF: Quad band pass filter, and M: mirror. **b**, Fluorescence micropipette for measuring 2D binding kinetics and real-time Ca^2+^ signaling. **c**, Cartoon illustration of 2D kinetic measurements of TCR-pMHC interactions at the cell membrane.

**Extended Data Fig. 3.**
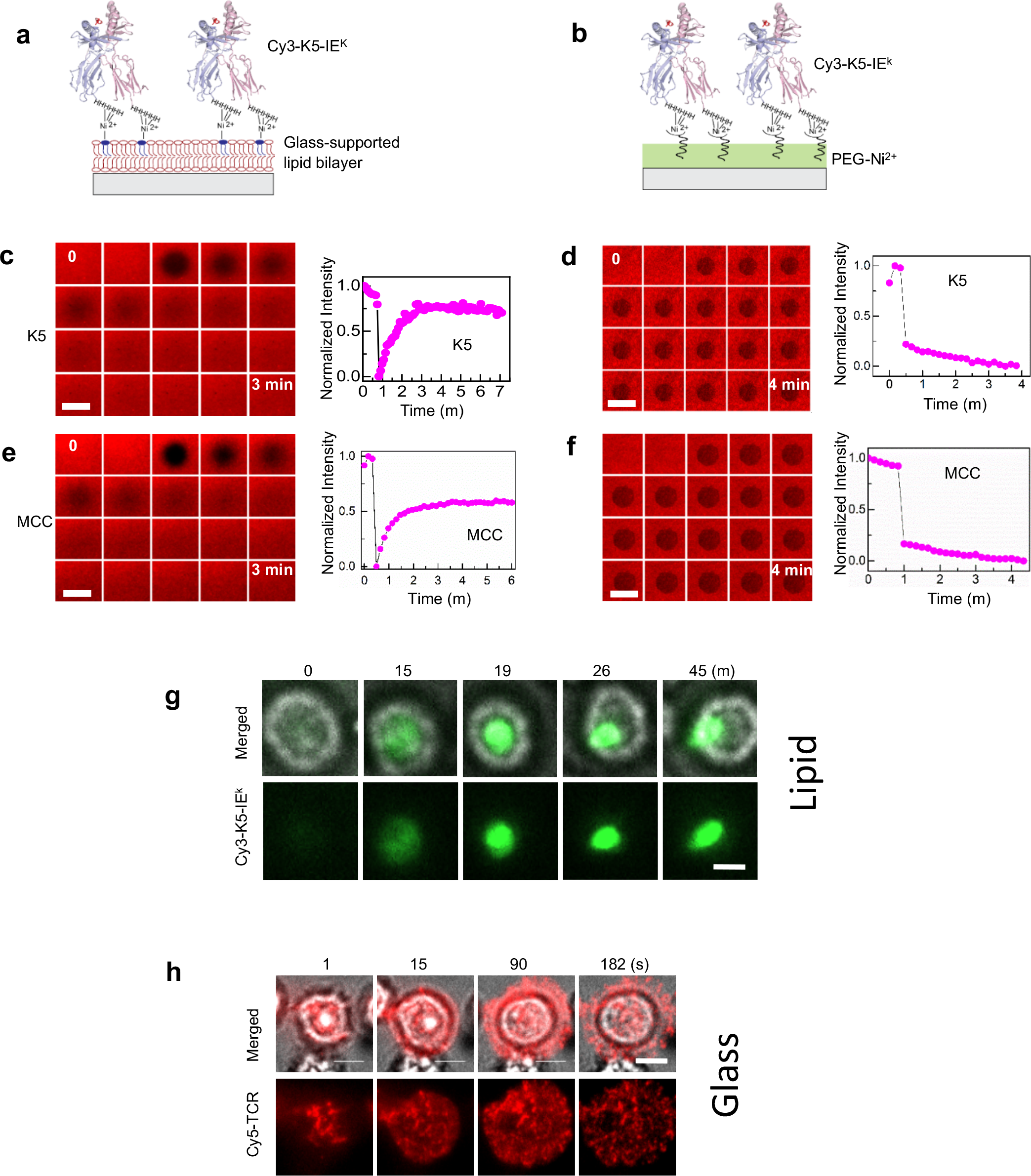
Functionalized lipid bilayer and glass surface. **a-b**, Cy3-labeled His_12_-K5-IE^k^ pMHCs were attached to Ni^2+^-functionalized glass supported lipid bilayer (a) and PEG-Ni^2+^ modified glass surface (b). **c-f**, Representative real-time fluorescence recovery after photobleaching (FRAP) images and corresponding time trajectories of fluorescent intensity for Cy3-lablled His_12_-K5-IE^k^ and His_12_-MCC-IE^k^ pMHCs attached on the lipid bilayer (c and e) and the PEG-Ni^2+^ modified glass surface (d and f). Three independent experiments were repeated for each pMHC at 37 °C. Fluorescence recovery confirmed the fluidity and integrity of lipid bilayer. No recovery was observed on glass surface. Also see Supplementary Movie 2. Scale bar: 10 μm. **g-h**, Immunological synapse formation on the lipid bilayer (g) and TCR microcluster formation on the PEG-Ni^2+^ glass surface (h). TCRs were labelled with Cy5 and pMHCs were labelled with Cy3. Also see Supplementary Movie 3.

**Extended Data Fig. 4.**
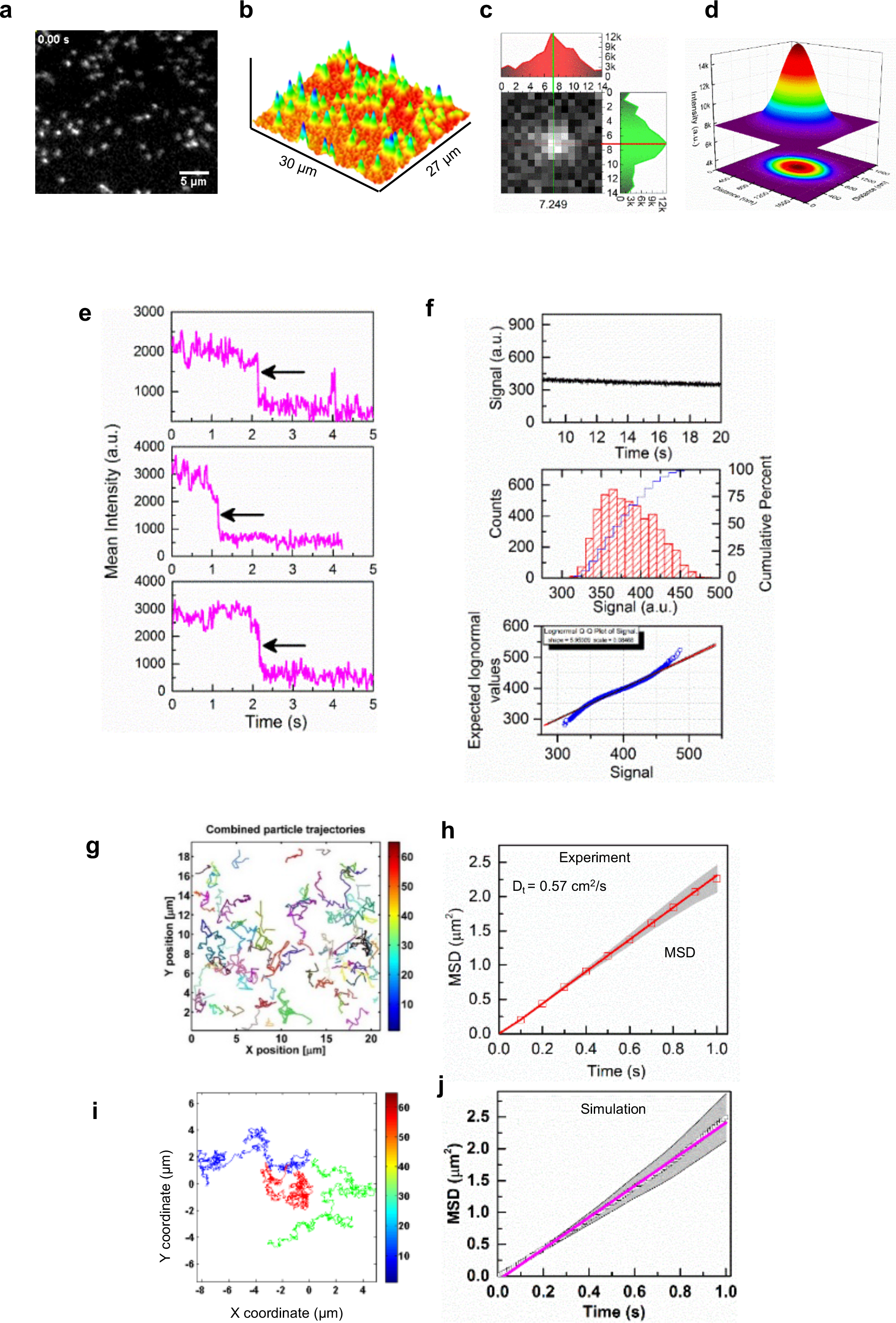
Single-molecule quantification of IE^K^ pMHCs on lipid bilayer. **a**, A representative TIRF image of single Cy3-labelled His_12_-K5-IE^k^ pMHCs on mobile lipid bilayer. Each white spot represents a single fluorescent pMHC. **b**, 3D fluorescence intensity profiles of single pMHCs shown in a. **c**, A representative 2D intensity profile of a single pMHC. **d**, The same molecule in c was fitted with a 3D Gaussian function to determine the full width at half maximum (FWHM), a value of 263 nm. This was comparable to the diffraction limit spot size (187 nm) for the 532-nm excitation laser. **e**, Single-step photobleaching confirmed the detection of single pMHCs on lipid bilayer. Representative single-step bleaching of three single Cy3-labelled His_12_-K5-IE^k^ pMHC molecules. **f**, The stability test of the 532-nm laser by examining the intensity profile of a standard fluorescence bead under continuous illumination. **g-h**, The diffusion of single pMHCs on lipid bilayer was tracked by a Fiji plugin TrackMate software (g) and the diffusion was quantified by a mean-square distance (MSD) plot (h). Also see Supplementary Movie 1. **i-j**, In silico simulated pMHC diffusion on lipid bilayer using a TrackArt program and the resulted MSD plot. The diffusion coefficients obtained from experiments and simulation were 0.57 ± 0.06 and 0.59 ± 0.09 μm/s, respectively.

**Extended Data Fig. 5.**
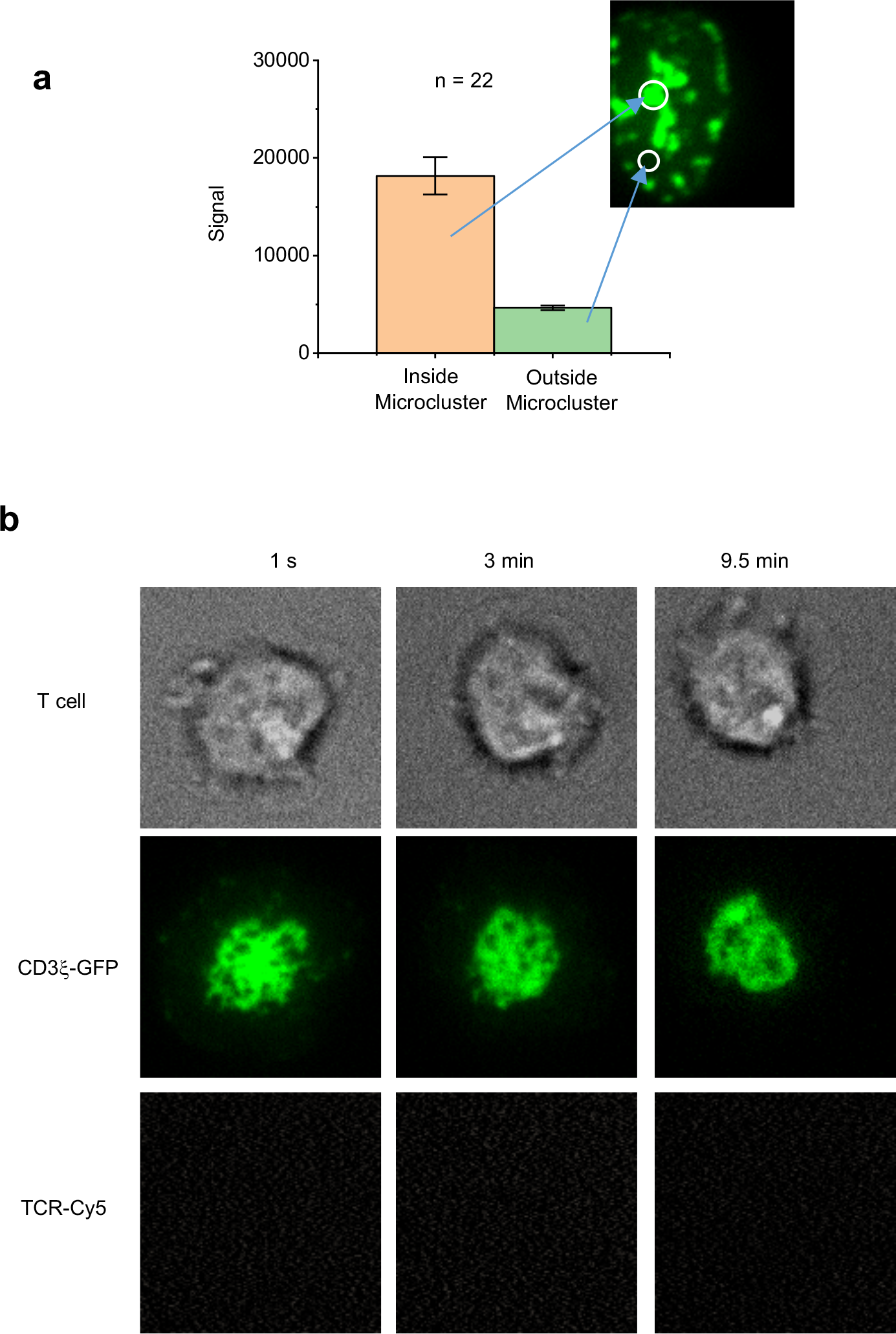
Transmembrane FRET is specific to GFP/Alexa568 FRET pair. **a**, Microclusters are signaling hotspot. CD3ζ-GFP signals inside and outside of the microclusters. **b**, No FRET is detected when Alexa568 is replaced by Cy5. A 470-nm LED light was used to excite GFP and measure GFP/Alexa568 FRET.

**Extended Data Fig. 6.**
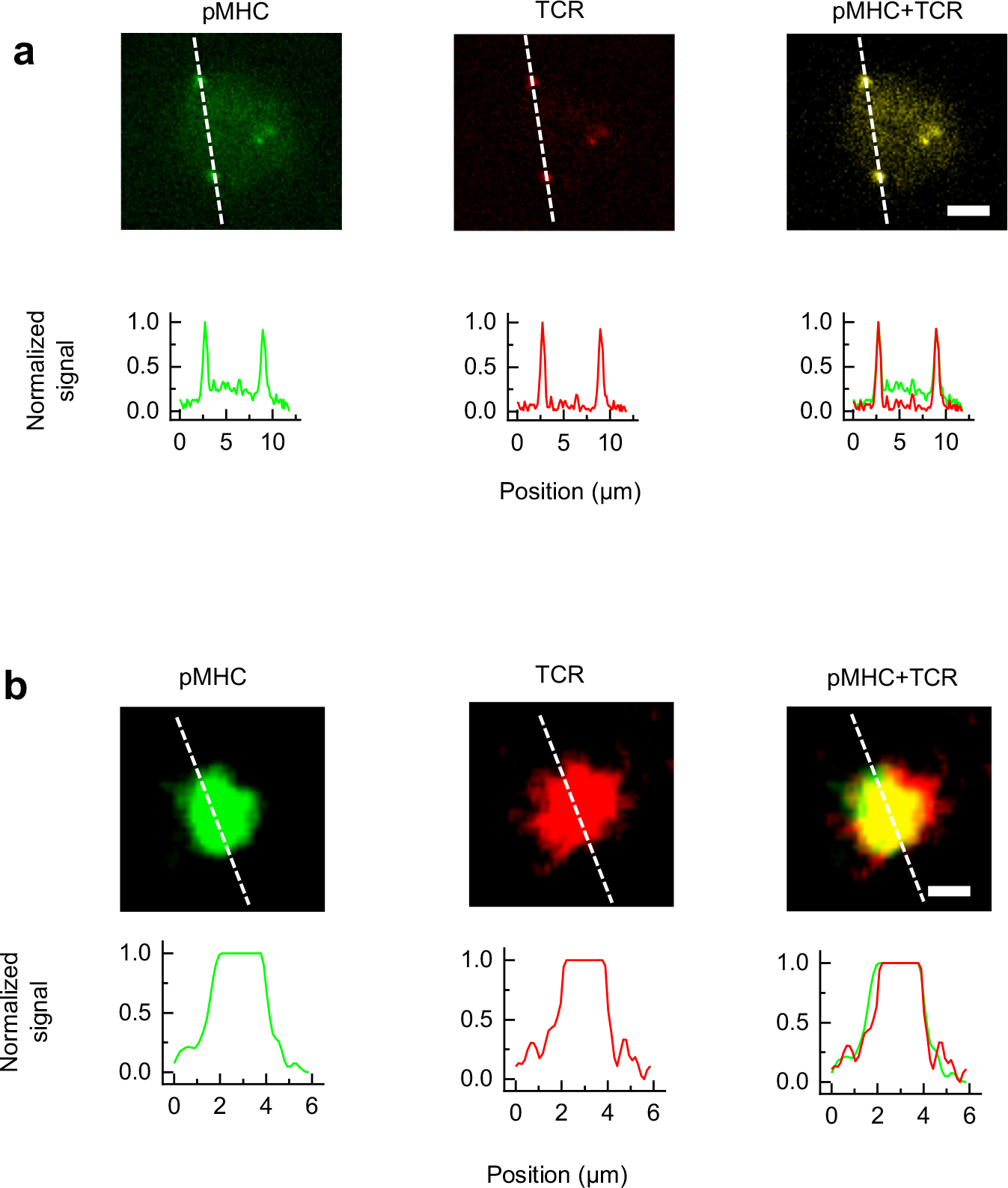
Co-localization between TCRs and pMHCs on lipid bilayer. Representative co-localization between TCR(s) and pMHC(s) in single-molecule **(a)** and ensemble **(b)** FRET experiments. Line scan was used to indicate the fluorescence profile and co-localization for both single-molecule **(a)** and ensemble **(b)** FRET. Scale bar is 5 μm.

**Extended Data Fig. 7.**
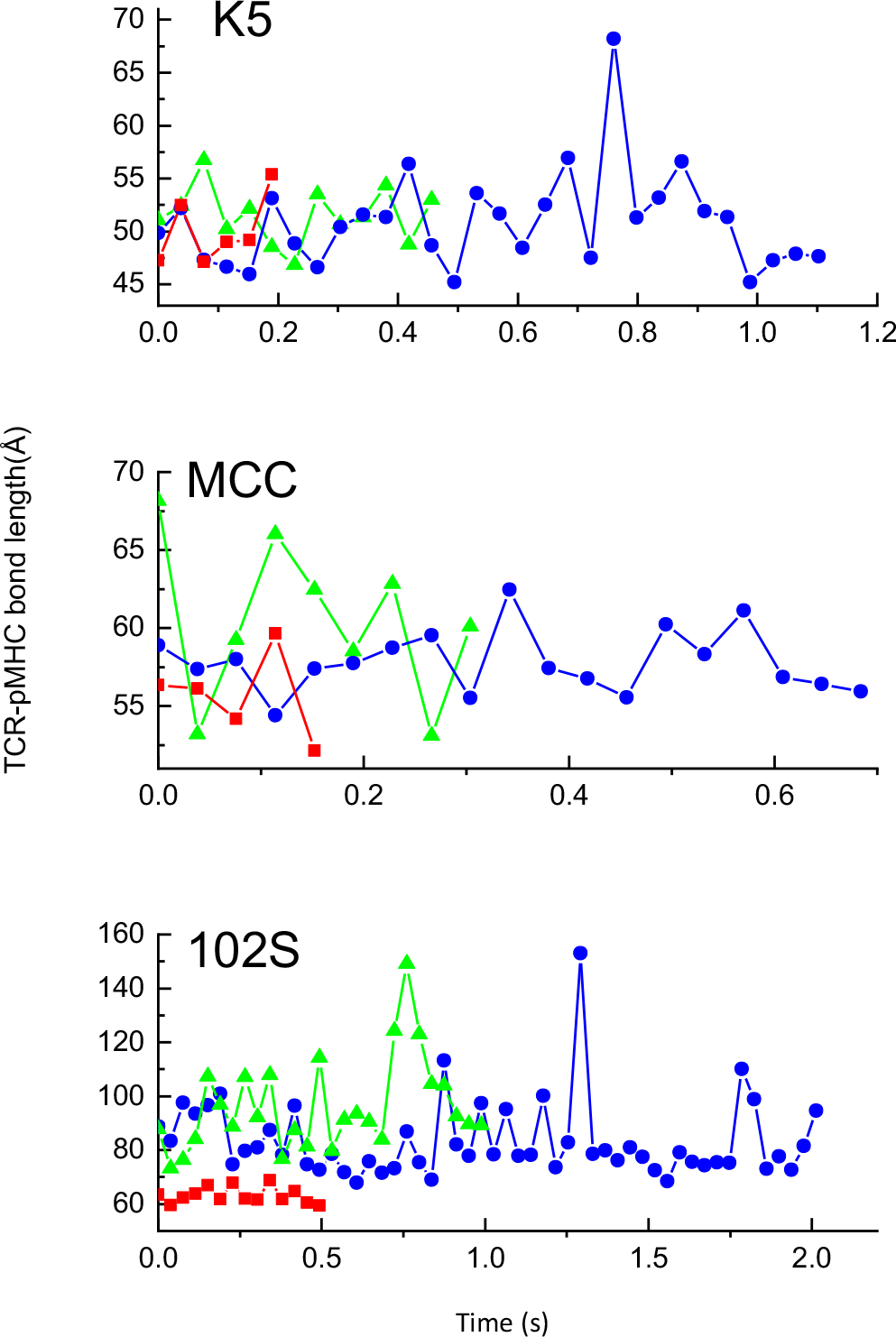
Representative bond length trajectories for K5, MCC and 102S pMHCs. For each peptide, three trajectories with long (blue), medium (green) and short (red) bond lifetimes were randomly chosen as demonstration. Note: the x- and y-axes are different for each peptide.

**Extended Data Fig. 8.**
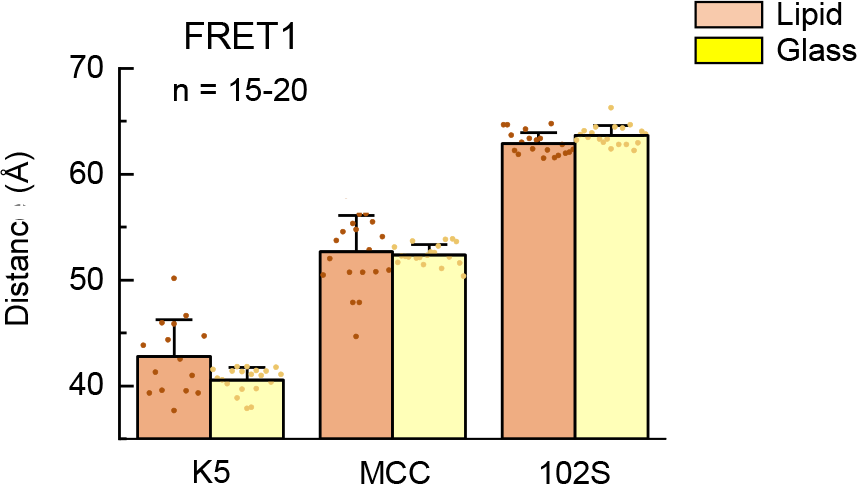
TCR-pMHC bond length measurements by ensemble FRET. The average bond lengths for K5, MCC and 102S pMHCs after ligation with 5C.C7 TCRs on a lipid bilayer and a glass surface.

**Extended Data Fig. 9.**
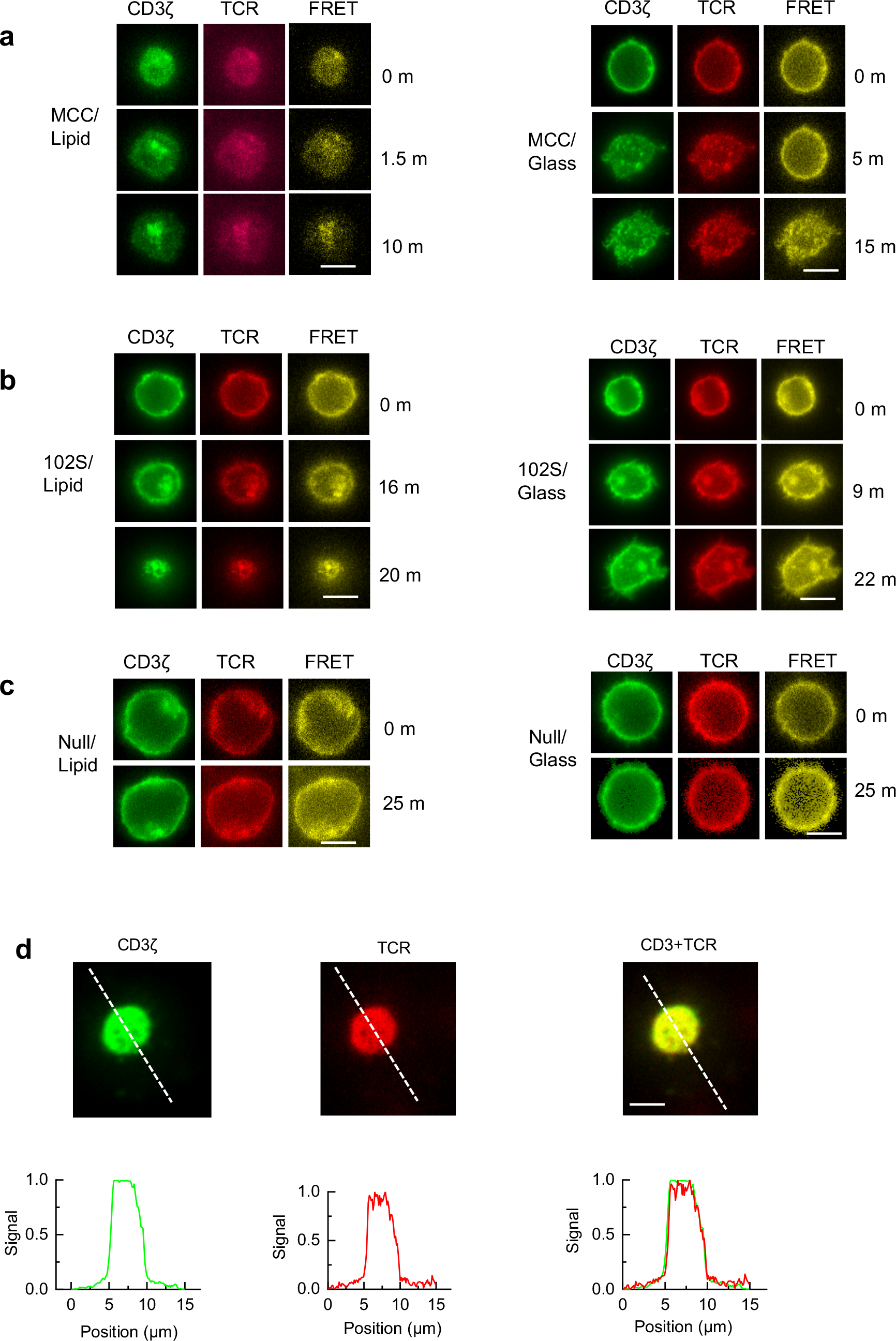
Transmembrane FRET measurements for MCC, 102S and Null pMHCs. **a-c**, Representative transmembrane FRET measurements between CD3ζ-GFP and TCR-Alexa568 on lipid bilayer (left) and glass surface (right) for MCC **(a)**, 102S **(b)** and Null **(c)**. Scale bar is 7 μm. **d**, Line scanning of CD3ζ, TCR and FRET channels for K5 pMHC after forming a stable immunological synapse on lipid bilayer. Scale bar is 5 μm.

**Extended Data Fig. 10.**
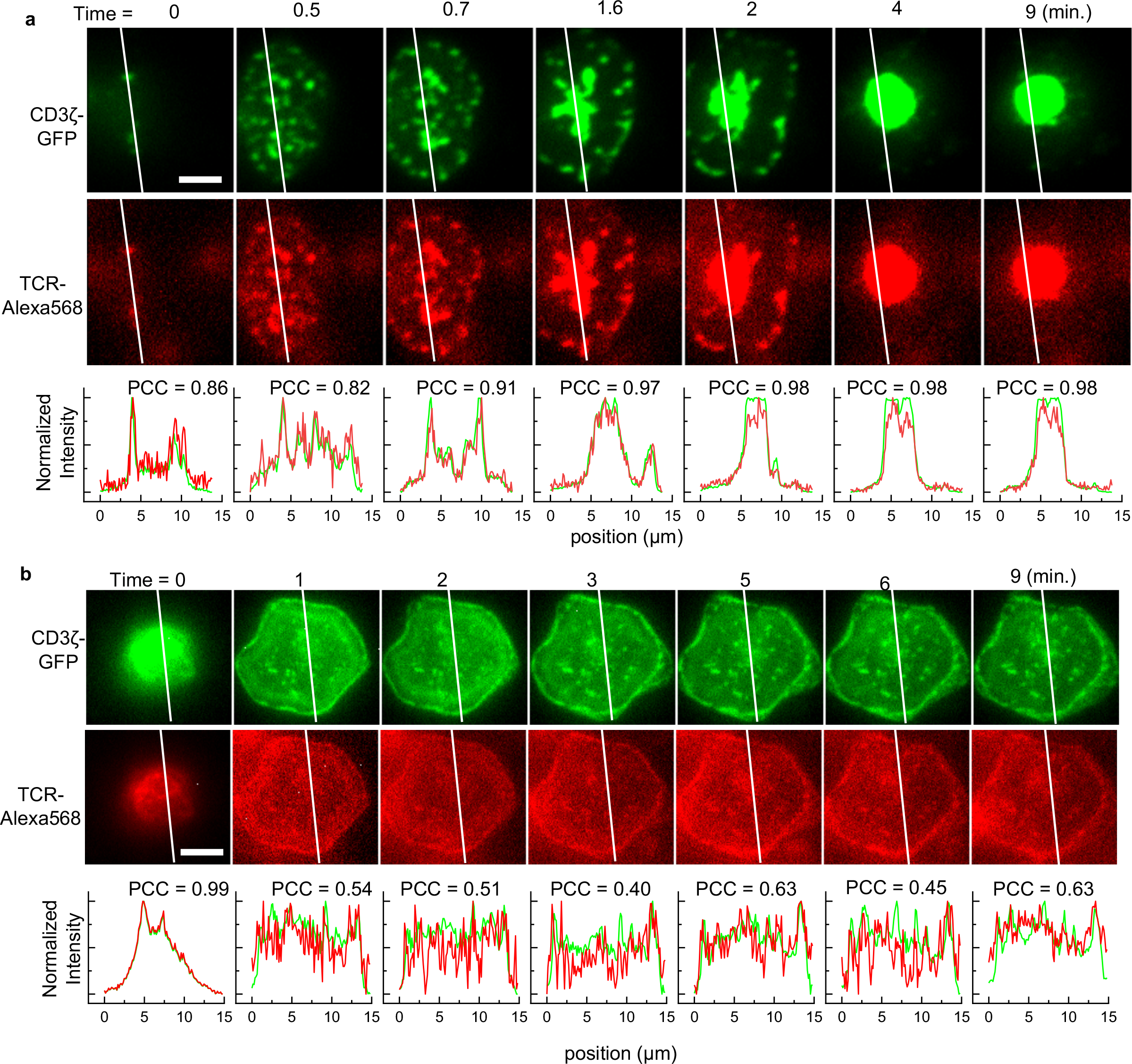
Colocalization of TCR and CD3ζ on lipid bilayer and glass surface. Time series of line scanning intensity profile for TCR and CD3ζ on lipid bilayer **(a)** and glass surface **(b)**. The colocalization between TCR and CD3ζ was quantified by Pearson correlation coefficient (PCC). The average PCC values are 0.93±0.07 and 0.59±0.19 on lipid bilayer and glass surface, respectively. Scale bar is 5 μm.

**Extended Data Fig. 11.**
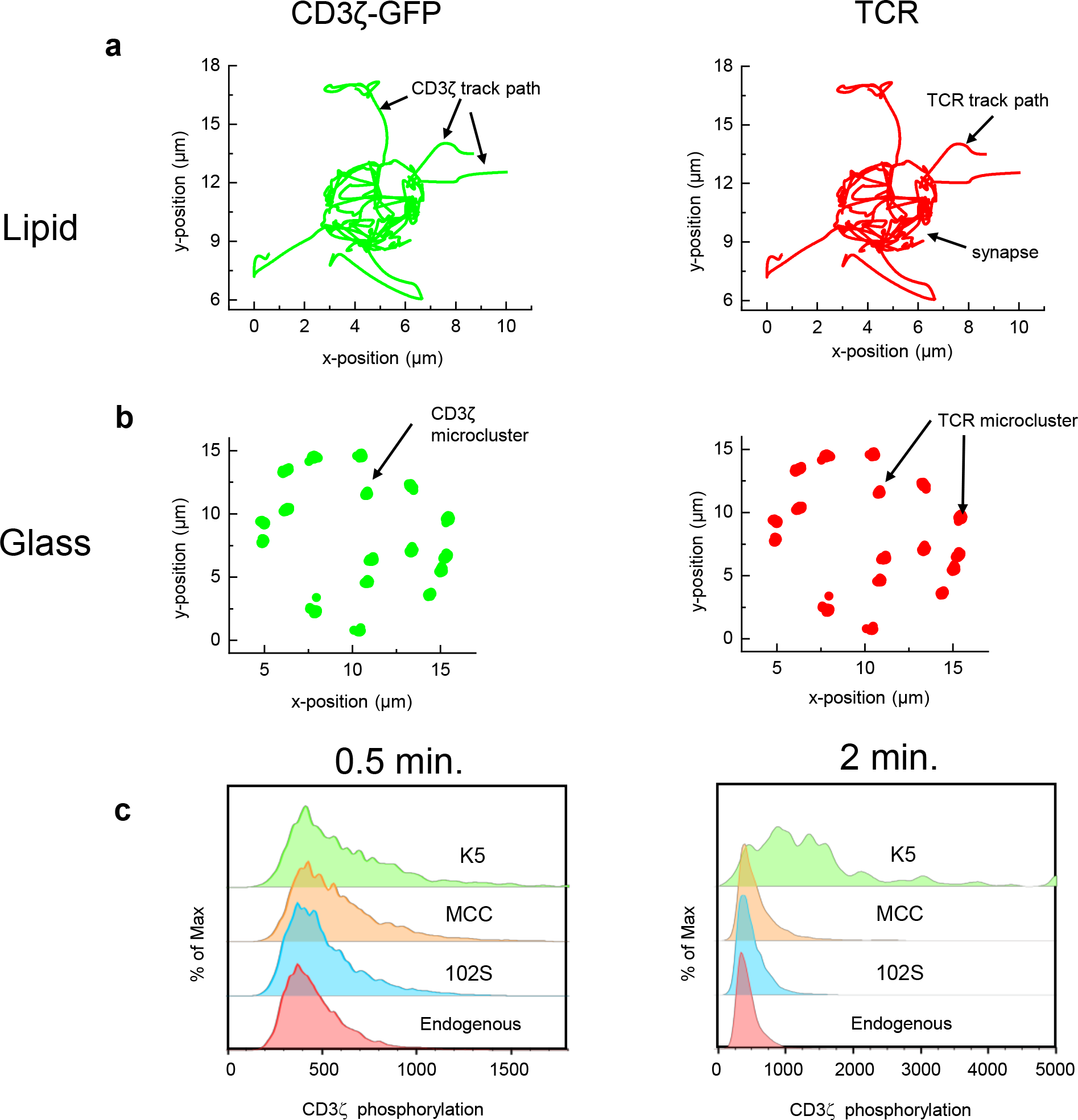
2D tracking of TCR and CD3ζ microclusters. **a-b**, Real-time tracking of CD3ζ (green) and TCR (red) microclusters on lipid bilayer **(a)** and glass surface **(b)** during transmembrane FRET experiments. Also see Supplementary Movie 5-7. **c**, Phospho-flow showing the phosphorylation of CD3ζ in T cells upon contacting antigen-presenting cells loaded with 102S, MCC or K5 peptides for 0.5 and 2 mins.

**Extended Data Fig. 12.**
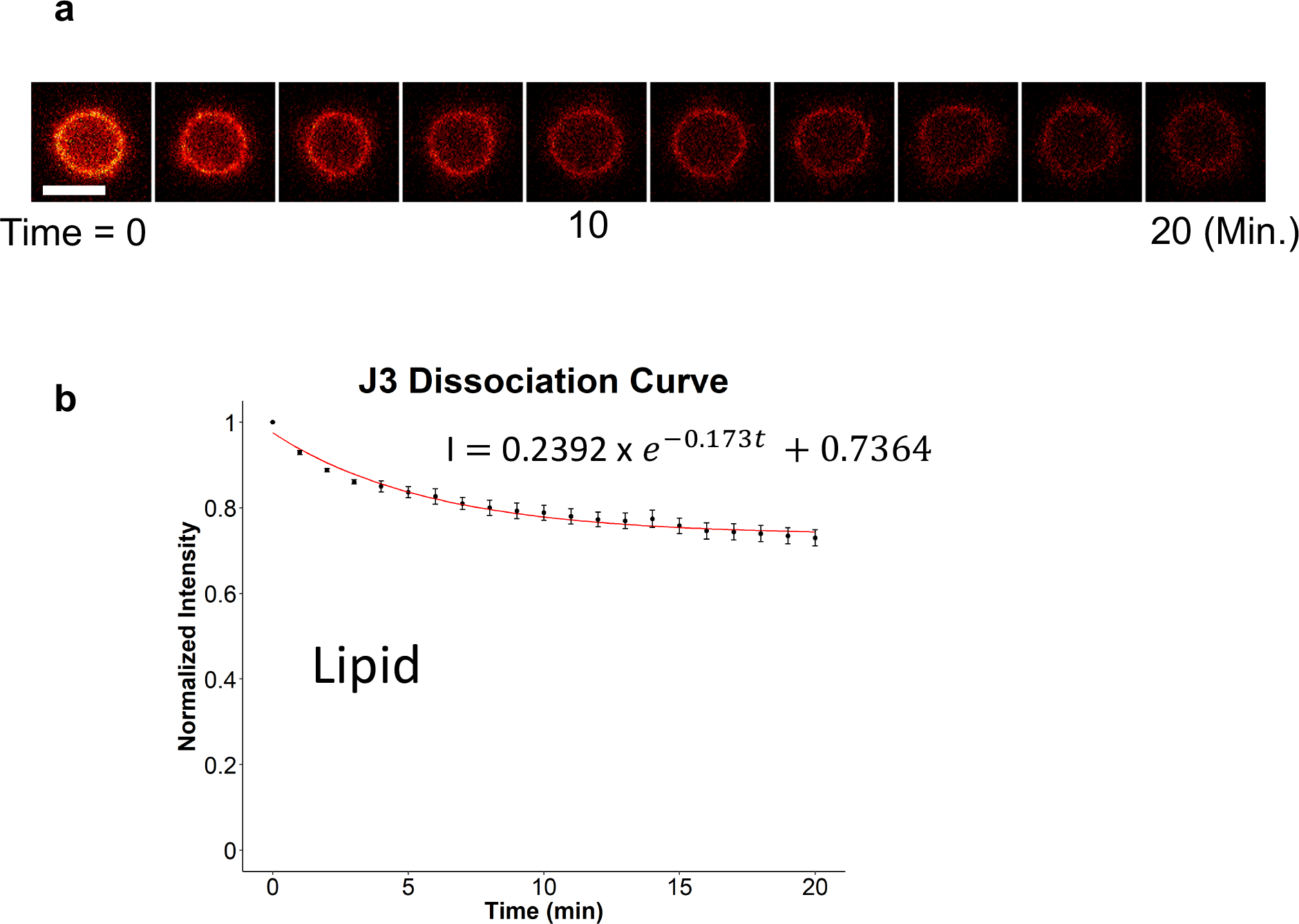
scFv disassociation calibration for transmembrane FRET. **a**, A representative time-lapse images of a J3 scFv labelled T cell. Scale bar is 10 μm. Fluorescence intensity is decreasing with time because of the dissociation of scFv from TCR. **b**, The intensity time trajectory was fitted with a first order disassociation model. Each data point was the average value of 50 independent measurements. The J3-Alexa568 intensities in transmembrane FRET experiments were corrected based on the dissociation kinetics of J3 scFv.

**Extended Data Fig. 13.**
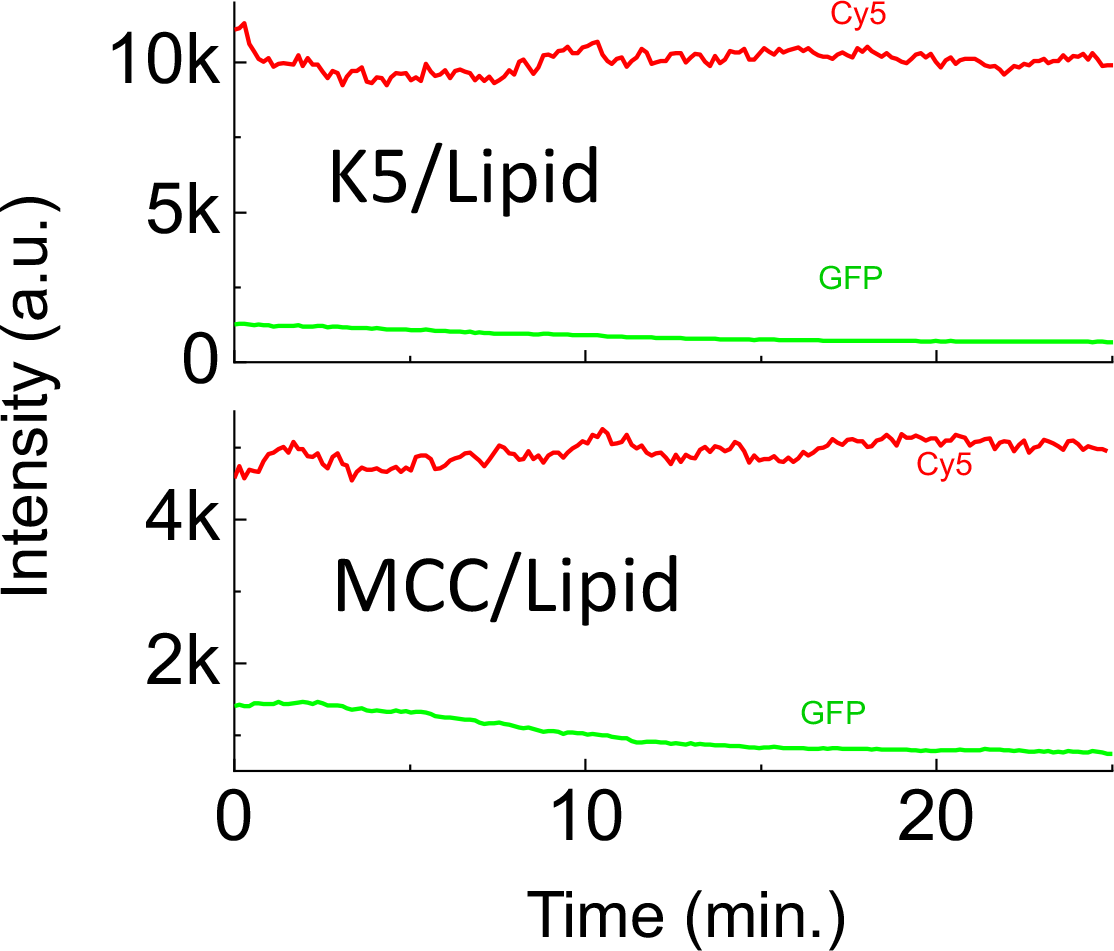
Negative controls of transmembrane FRET. To rule out the possibility that transmembrane FRET in Fig. 3 was caused by different diffusion rates of TCRs and CD3ζ into a microcluster due to the ratiometric nature of FRET, we labelled TCRs with Cy5 and CD3ζ with GFP (GFP/Cy5 is a non-FRET pair). The GFP was excited by a 470-nm LED light and the Cy5 was excited by a 640-nm LED light sequentially. The mean fluorescence intensities of Cy5 and GFP (and their ratios) stayed constant in a representative microcluster upon TCR-pMHC ligation for K5 and MCC pMHCs. These experiments confirmed that the transmembrane FRET between CD3ζ-GFP and TCR-Alexa568 in Fig. 3 were merely due to TCR-CD3ζ conformational changes, as well as confirmed that TCR-CD3ζ complexes are very stable (also see TCR/CD3 colocalization in Extended Data Fig. 10a). The mean fluorescence intensities were recorded in real time upon adding fluorescently labelled T cells to K5 (top panel) or MCC (bottom panel) pMHCs containing lipid bilayer. Shown are representative time trajectories of intensity out of 3 independent experiments for each pMHC. Note: these experiments were different from those shown in Extended Data Fig. 5B, which only used a single 470-nm LED light to excite GFP.

**Extended Data Fig. 14.**
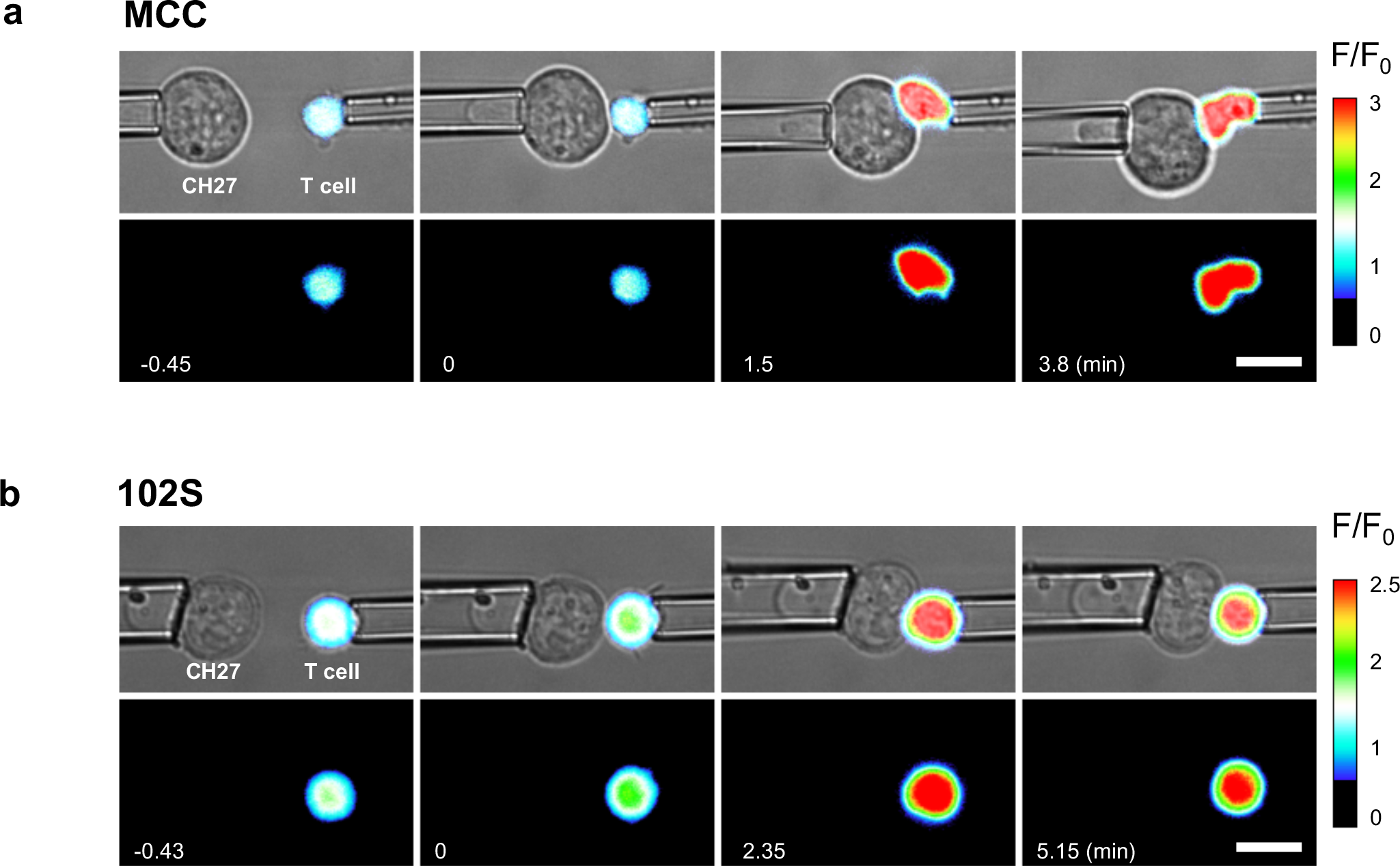
Real-time Ca^2+^ signaling at the single-cell level. Calcium imaging of a T cell contacting a CH27 cell loaded with MCC (**a**) or 102S (**b**) peptide measured by fluorescent micropipette at 37 °C. Intracellular high Ca^2+^ concentration is indicated by the hot color. Fluorescence signal was recorded by time-lapse fluorescence microscopy (Extended Data Fig. 2b) and the fold-increase of Ca^2+^ signaling (F/F_0_) was shown by pseudo color. Shown are representative experiments out of 3-5 independent experiments for each peptide (MCC and 102S). See Fig. 4e-f for Ca^2+^ signaling of T cell activated by K5 and Null peptides. Also see Supplementary Movie 9. Scale bar 10 μm.

**Extended Data Fig. 15.**
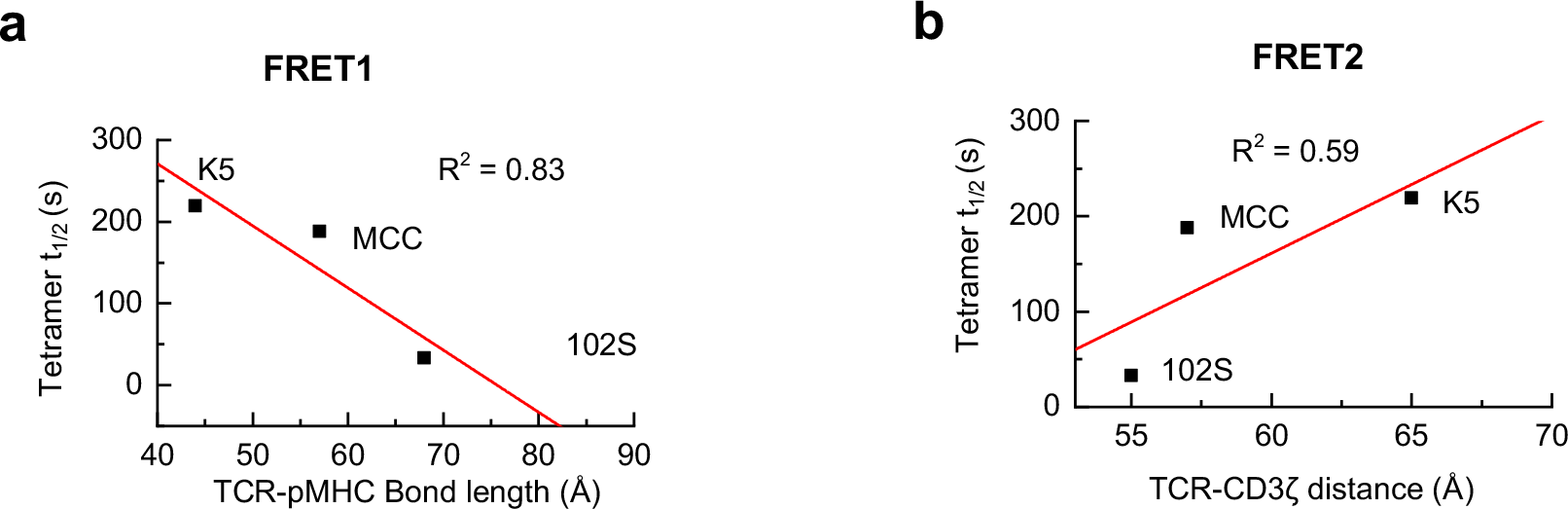
Correlation between of the half-lives (t_1/2_) of tetramer staining at 25 °C and intermolecular TCR-pMHC bond length (a) or intramolecular TCR-CD3ζ distance (b). The tetramer t_1/2_ data was adopted from Corse et al. 2010.

**Extended Data Table 1.**
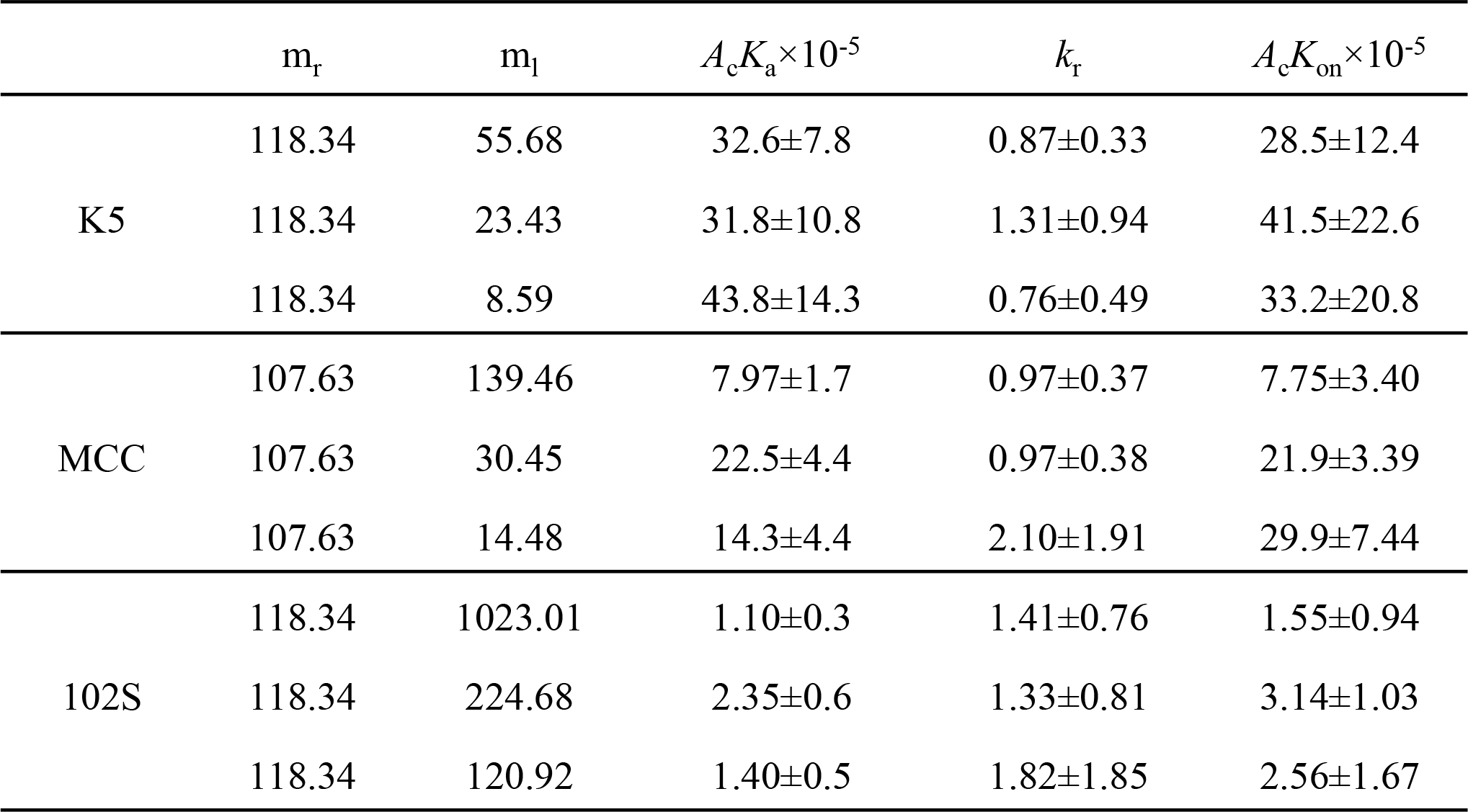
2D kinetic parameters.

## References

1. Irvine, D. J., Purbhoo, M. A., Krogsgaard, M. & Davis, M. M. Direct observation of ligand recognition by T cells. Nature 419, 845–849 (2002).

2. Huang, J. et al. A Single peptide-major histocompatibility complex ligand triggers digital cytokine secretion in CD4+ T Cells. Immunity 39, 846–857 (2013).

3. Huppa, J. B. et al. TCR-peptide-MHC interactions in situ show accelerated kinetics and increased affinity. Nature 463, 963–967 (2010).

4. Huang, J. et al. The kinetics of two-dimensional TCR and pMHC interactions determine T-cell responsiveness. Nature 464, 932–936 (2010).

5. Ha, T. et al. Probing the interaction between two single molecules: fluorescence resonance energy transfer between a single donor and a single acceptor. Proc. Natl. Acad. Sci. 93, 6264–6268 (1996).

6. van der Merwe, P. A. & Dushek, O. Mechanisms for T cell receptor triggering. Nat Rev Immunol 11, 47–55 (2010).

7. Rudolph, M. G., Stanfield, R. L. & Wilson, I. A. How TCRs bind MHCs, peptides, and coreceptors. Annu. Rev. Immunol. 24, 419–66 (2006).

8. Sasmal, D. K., Pulido, L. E., Kasal, S. & Huang, J. Single-molecule fluorescence resonance energy transfer in molecular biology. Nanoscale 8, 19928–19944 (2016).

9. Corse, E., Gottschalk, R. A., Krogsgaard, M. & Allison, J. P. Attenuated T Cell Responses to a High-Potency Ligand In Vivo. PLoS Biol. 8, e1000481 (2010).

10. Natarajan, K. et al. An allosteric site in the T-cell receptor Cβ domain plays a critical signalling role. Nat. Commun. 8, 15260–13 (2017).

11. Newell, E. W. et al. Structural Basis of Specificity and Cross-Reactivity in T Cell Receptors Specific for Cytochrome c-I-E. J. Immunol. 186, 5823–5832 (2011).

12. Yang, H. et al. Protein conformational dynamics probed by single-molecule electron transfer. Science 302, 262–6 (2003).

13. Choudhuri, K., Wiseman, D., Brown, M. H., Gould, K. & van der Merwe, P. A. T-cell receptor triggering is critically dependent on the dimensions of its peptide-MHC ligand. Nature 436, 578–582 (2005).

14. Choudhuri, K. et al. Peptide-Major Histocompatibility Complex Dimensions Control Proximal Kinase-Phosphatase Balance during T Cell Activation. J. Biol. Chem. 284, 26096–26105 (2009).

15. Hui, E. et al. T cell costimulatory receptor CD28 is a primary target for PD-1-mediated inhibition. Science 355, 1428–1433 (2017).

16. Davis, S. J. & van der Merwe, P. A. The kinetic-segregation model: TCR triggering and beyond. Nat. Immunol. 7, 803–809 (2006).

17. Mossman, K. D., Campi, G., Groves, J. T. & Dustin, M. L. Altered TCR signaling from geometrically repatterned immunological synapses. Science 310, 1191–3 (2005).

18. Yokosuka, T. et al. Newly generated T cell receptor microclusters initiate and sustain T cell activation by recruitment of Zap70 and SLP-76. Nat Immunol 6, 1253–1262 (2005).

19. Campi, G., Varma, R. & Dustin, M. L. Actin and agonist MHC-peptide complex-dependent T cell receptor microclusters as scaffolds for signaling. J. Exp. Med. 202, 1031–6 (2005).

20. Taylor, M. J., Husain, K., Gartner, Z. J., Mayor, S. & Vale, R. D. A DNA-Based T Cell Receptor Reveals a Role for Receptor Clustering in Ligand Discrimination. Cell 169, 108–119.e20 (2017).

21. Berg, J. M., Tymoczko, J. L., Stryer, L. & Stryer, L. Biochemistry, 5th Ed. (W H Freeman, New York).

22. Liu, B., Chen, W., Evavold, B. D. & Zhu, C. Accumulation of dynamic catch bonds between TCR and agonist peptide-MHC triggers T cell signaling. Cell 157, 357–368 (2014).

23. Basu, R. et al. Cytotoxic T Cells Use Mechanical Force to Potentiate Target Cell Killing. Cell 165, 100–110 (2016).

24. Sibener, L. V et al. Isolation of a Structural Mechanism for Uncoupling T Cell Receptor Signaling from Peptide-MHC Binding. Cell 174, 672–687.e27 (2018).

25. Shi, X. et al. Ca2+regulates T-cell receptor activation by modulating the charge property of lipids. Nature 493, 111–115 (2013).

26. Chakraborty, A. K. & Weiss, A. Insights into the initiation of TCR signaling. Nat. Immunol. 15, 798–807 (2014).

27. Gil, D., Schamel, W. W. A., Montoya, M., Sánchez-Madrid, F. & Alarcón, B. Recruitment of Nck by CD3ϵ Reveals a Ligand-Induced Conformational Change Essential for T Cell Receptor Signaling and Synapse Formation. Cell 109, 901–912 (2002).

28. Xu, C. et al. Regulation of T Cell Receptor Activation by Dynamic Membrane Binding of the CD3ɛ Cytoplasmic Tyrosine-Based Motif. Cell 135, 702–713 (2008).

29. Guo, X. et al. Lipid-dependent conformational dynamics underlie the functional versatility of T-cell receptor. Cell Res. 27, 505–525 (2017).

30. Lee, M. S. et al. A Mechanical Switch Couples T Cell Receptor Triggering to the Cytoplasmic Juxtamembrane Regions of CD3ζζ. Immunity 43, 227–239 (2015).

## References (methods)

31. Birnbaum, M. E. et al. Deconstructing the peptide-MHC specificity of t cell recognition. Cell 157, 1073–1087 (2014).

32. Howarth, M. & Ting, A. Y. Imaging proteins in live mammalian cells with biotin ligase and monovalent streptavidin. Nat. Protoc. 3, 534–545 (2008).

33. Tsumoto, K. et al. Highly efficient recovery of functional single-chain Fv fragments from inclusion bodies overexpressed in Escherichia coli by controlled introduction of oxidizing reagent - Application to a human single-chain Fv fragment. J. Immunol. Methods 219, 119–129 (1998).

34. Ozaki, C. Y. et al. Single Chain Variable Fragments Produced in Escherichia coli against Heat-Labile and Heat-Stable Toxins from Enterotoxigenic E. coli. PLoS One 10, e0131484 (2015).

35. Leisegang, M. et al. Eradication of large solid tumors by gene therapy with a T-cell receptor targeting a single cancer-specific point mutation. Clin. Cancer Res. 22, 2734–2743 (2016).

36. Su, X. et al. Phase separation of signaling molecules promotes T cell receptor signal transduction. Science 352, 595–9 (2016).

37. Matysik, A. & Kraut, R. S. TrackArt: the user friendly interface for single molecule tracking data analysis and simulation applied to complex diffusion in mica supported lipid bilayers. BMC Res. Notes 7, 274 (2014).

38. Edelstein, A. D. et al. Advanced methods of microscope control using μManager software. J. Biol. Methods 1, 10 (2014).

39. Huang, J., Edwards, L. J., Evavold, B. D. & Zhu, C. Kinetics of MHC-CD8 Interaction at the T Cell Membrane. J. Immunol. 179, 7653–7662 (2007).

40. Roy, R., Hohng, S. & Ha, T. A practical guide to single-molecule FRET. Nat. Methods 5, 507–516 (2008).

41. Chandler, D. Introduction to Modern Statistical Mechanics. Oxford University Press (1987).

42. Tinevez, J. Y. et al. TrackMate: An open and extensible platform for single-particle tracking. Methods 115, 80–90 (2017).

43. Krutzik, P. O., Trejo, A., Schulz, K. R. & Nolan, G. P. Phospho Flow Cytometry Methods for the Analysis of Kinase Signaling in Cell Lines and Primary Human Blood Samples. in Methods in molecular biology (Clifton, N.J.) 699, 179–202 (2011).

44. Dominguez, D. et al. Exogenous IL-33 Restores Dendritic Cell Activation and Maturation in Established Cancer. J. Immunol. 198, 1365–1375 (2017).

45. Huse, M. et al. Spatial and Temporal Dynamics of T Cell Receptor Signaling with a Photoactivatable Agonist. Immunity 27, 76–88 (2007).

